# Negative emotional behavior during fentanyl abstinence is mediated by adaptations in nucleus accumbens neuron subtypes

**DOI:** 10.1101/2022.05.15.491856

**Authors:** Megan E. Fox, Andreas B. Wulff, Daniela Franco, Eric Choi, Cali A. Calarco, Michel Engeln, Makeda D. Turner, Ramesh Chandra, Victoria M. Rhodes, Scott M. Thompson, Seth A. Ament, Mary Kay Lobo

**Affiliations:** Department of Anesthesiology, Department of Pharmacology, Penn State College of Medicine, Hershey, PA, USA; Department of Anatomy & Neurobiology, University of Maryland School of Medicine, Baltimore, MD, USA; Department of Physiology, University of Maryland School of Medicine, Baltimore, MD, USA; Institute for Genome Sciences, University of Maryland School of Medicine, Baltimore, MD, USA; Department of Psychiatry, University of Maryland School of Medicine, Baltimore, MD, USA; University of Bordeaux, CNRS, INCIA, UMR 5287, F-33000 Bordeaux, France

**Keywords:** Opioids, nucleus accumbens, abstinence, E2f1, medium spiny neurons, fentanyl

## Abstract

Opioid discontinuation generates a withdrawal syndrome marked by a negative emotional state. Increased anxiety and dysphoria during opioid discontinuation are a significant barrier to achieving long-term abstinence in opioid-dependent individuals. Adaptations in brain-reward circuitry are implicated in the opioid abstinence syndrome, but current knowledge is limited to changes following natural and semi-synthetic opioids. Here we report abstinence from the synthetic opioid fentanyl engenders structural, functional, and molecular plasticity in nucleus accumbens neuron subtypes (MSNs) that mediate negative emotional behaviors. We show fentanyl abstinence causes dendritic atrophy and increased excitatory drive exclusive to D1-receptor containing MSNs. Using subtype specific RNAseq and Weighted Gene Co-Expression Network Analysis, we identified molecular signatures of fentanyl abstinence in MSN subtypes. We found a network of co-expressed genes downregulated selectively in D1-MSNs, and transcriptionally co-regulated by E2F1. We show targeting abstinence-induced molecular changes protects D1-MSNs from maladaptive plasticity and alleviates negative emotional behaviors after fentanyl abstinence.

## Introduction

Rates of opioid misuse, relapse, and deadly overdose have recently skyrocketed in North America. Drug seizure data indicates that many street drugs and counterfeit opioid pills contain significant levels of the synthetic opioid fentanyl, and that these drugs are often sold to unsuspecting users (Drug Enforcement Administration, 2016). Highly potent synthetic opioids accounted for 75% of overdose deaths in 2020 (Ahmad et al., 2021), but remain broadly understudied. Generally, opioid exposure and abstinence engage nuclei within the brain’s reward circuitry and cause lasting molecular changes thought to alter circuit function and promote persistent drug use and relapse (Volkow et al., 2016). One hallmark of opioid use is increased negative affect during withdrawal and abstinence (Koob, 2020). Indeed, managing these symptoms is an important component of medication assisted therapies for opioid use disorder, yet our mechanistic understanding of how opioid abstinence changes the brain is primarily derived from studies using natural or semi-synthetic opioids (See (Browne et al., 2020; Jordan and Xi, 2022; Reiner et al., 2019) for recent reviews).

The nucleus accumbens (NAc) is a key locus in brain reward-circuitry that undergoes distinct molecular, cellular, and structural changes during opioid abstinence. Opioid withdrawal reduces dopamine release in the NAc (Acquas et al., 1991; Fox et al., 2017; Pothos et al., 1991; Rossetti et al., 1992) and induces numerous transcriptional changes dependent on opioid, sex, and abstinence duration (Cahill et al., 2018; Ferguson et al., 2013; Hofford et al., 2021; Martin et al., 2019; Mayberry et al., 2022; Spijker et al., 2004; Sun et al., 2016; Townsend et al., 2021). Opioid cessation also causes morphological and electrophysiological changes to NAc neurons. Most studies show decreased dendritic spine density and increased excitatory drive, however the physiologic responses are heterogenous, often NAc subregion specific, and in some cases there are no changes or a decrease in excitability dependent on the paradigm (Diana et al., 2006; Graziane et al., 2016; Hearing et al., 2016; Heng et al., 2008; Madayag et al., 2019; Matsubara et al., 1999; McDevitt et al., 2019; Pal and Das, 2013; Robinson and Kolb, 2004; Spiga et al., 2005; Thompson et al., 2021; Wu et al., 2012, 2013).

The NAc, located in the ventral striatum, contains primarily medium sized GABAergic spiny projection neurons that are divided into two subtypes based on dopamine receptor expression. Drd1 or Drd2 expressing neurons (D1- and D2-MSNs, respectively) have little overlap in their projection targets (Gerfen and Surmeier, 2011; Kupchik et al., 2015; Smith et al., 2013), and often have opposing roles in driving drug- and reward-related behaviors (Calipari et al., 2016; Hauser et al., 2015; Hikida et al., 2010; James et al., 2013; Koo et al., 2014; Kravitz et al., 2012; Lobo et al., 2010; Tai et al., 2012). Increased activity in D1-MSNs is typically considered to be “pro-reward” and D2-MSNs “anti-reward,” however there are an increasing number of exceptions to this dichotomy in different parts of the striatum (Cui et al., 2013; Gallo et al., 2018; Gibson et al., 2018; Natsubori et al., 2017; Soares-Cunha et al., 2016, 2018, 2019; Vicente et al., 2016). Much less is known about specific MSN subtypes in opioid abstinence (Graziane et al., 2016; Hearing et al., 2016; Madayag et al., 2019; Martin et al., 2019; McDevitt et al., 2019), especially abstinence from synthetic opioids, and females were often excluded from previous studies. Further, many studies house animals singly, which is an added stressor that impacts drug-related behavior (Bozarth et al., 1989; Engeln et al., 2021; Kennedy et al., 2012; Westenbroek et al., 2013).

Based on our previous work showing NAc D1-MSNs drive stress-induced anhedonia and social avoidance (Fox et al., 2020a; Francis et al., 2017, 2019), we hypothesized D1-MSNs play a key role in the increased negative emotional behaviors during synthetic opioid abstinence. Here, we use numerous neuron subtype specific techniques to show fentanyl abstinence causes dendritic atrophy, increased excitatory drive, and molecular changes specific to NAc D1-MSNs in both sexes. We further show that negative-affective behaviors and NAc D1-MSN morphologic and physiologic changes can be prevented by targeting the transcriptional regulator E2f1.

## Results

### Homecage fentanyl abstinence is associated with increased negative emotional behavior and reduced dendritic complexity of D1-neurons

Due to the stress-sensitivity of NAc MSNs, we used a low-stress fentanyl exposure and abstinence paradigm that allowed for intermittent drug delivery and minimized handling and social isolation stress. We first established our homecage fentanyl drinking and abstinence paradigm was sufficient to increase negative-affect and stress-like behaviors typically associated with opioid abstinence in both sexes (Timeline in **Fig 1A**. Sexes combined due to no significant sex effects; **Supplemental File 1**). We found reduced social-interaction time in a 3-chamber social interaction test (**Fig 1B**, p=0.047) and decreased time in the open arms of an elevated plus maze in fentanyl abstinent mice (**Fig 1C**, p=0.019). To assess if fentanyl abstinence enhanced susceptibility to mild stress, we tested mice for social avoidance after an abbreviated social defeat stress (**Fig 1D**) and found increased stress-susceptibility (decreased interaction time) in fentanyl abstinent mice subject to brief social stress (**Fig 1E**, p=0.0017, Sidak’s post-hoc after 2-way ANOVA). To assess if the fentanyl paradigm was sufficient to generate physical dependence, we administered 1 mg/kg naloxone to fentanyl and water mice and observed precipitated withdrawal signs. We found increased global withdrawal scores in fentanyl-naloxone mice relative to water-naloxone (**Fig 1F**, p=0.002; fentanyl drinking data for all mice in **Fig S1A**).

**Figure 1.**
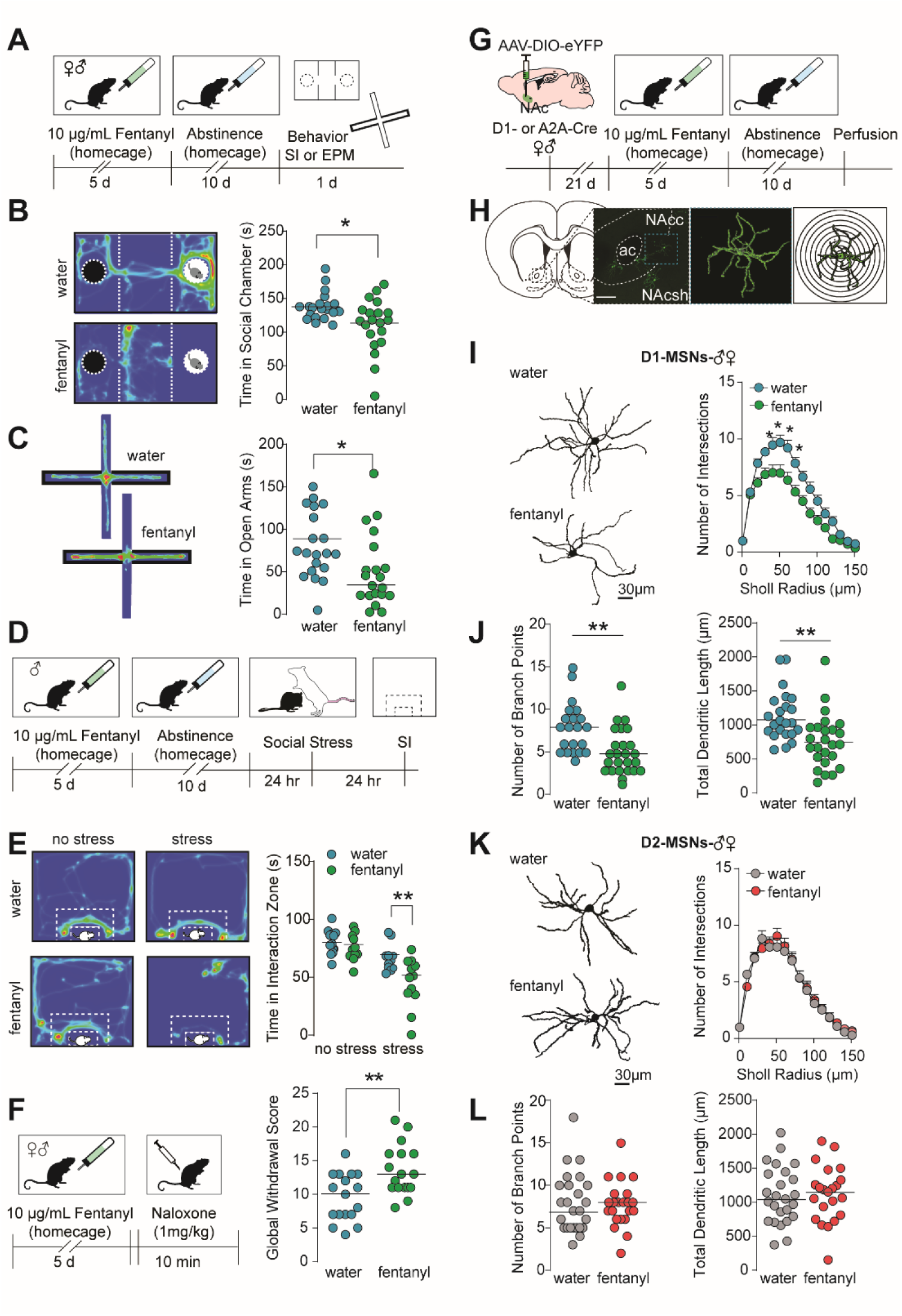
Fentanyl abstinence increases negative emotional behaviors and reduces dendritic complexity of D1-MSNs. **(A)** Timeline for homecage fentanyl abstinence paradigm. Male and female mice received 10µg/mL fentanyl in drinking water for 5 days, then underwent 10 days of abstinence. Mice then underwent either social interaction (SI) or elevated plus maze (EPM) testing. (**B**) *Left*: example heat maps showing time a water or fentanyl abstinent mouse spent in the 3-chamber social interaction chamber when a novel mouse was present in the rightmost chamber. Warmer colors indicate increased duration. *Right*: Median and individual data points showing time spent in the social chamber in water or fentanyl abstinent mice with a social target present (*, p=0.0165). (**C**) *Left*: example heat maps showing time a water or fentanyl abstinent mouse spent in the open and closed arms of an elevated plus maze. The closed arms are denoted by the thick black border. *Right*: Median and individual data points showing time spent in the open arms of the EPM (*, p=0.008). (**D**) Timeline for stress-susceptibility testing. After homecage fentanyl abstinence, male mice underwent a 1-day social stress paradigm in which they were subject to 3 brief agonistic encounters with 3 aggressive CD-1 residents separated by 10 min of sensory interaction. 24hr after stress, mice were assessed for social avoidance in an open field based social interaction (SI) test. (**E**) *Left*: example heat maps showing time unstressed and stressed water or fentanyl mice spent interacting with a novel CD-1. White dashed lines indicate interaction zone. *Right*: Median and individual data points showing time spent interacting with the social target (**, p=0.0017, Sidak’s post-hoc after 2-way ANOVA). (**F**) *Left*: timeline for naloxone precipitated withdrawal testing. Following 5 days of homecage fentanyl, male and female mice received 1 mg/kg naloxone and were observed for withdrawal signs. *Right*: Median and individual data points showing global withdrawal scores in water or fentanyl mice given naloxone (**, p=0.0021). (**G**) Schematic of sparse-labeling approach of D1- and D2-MSNs. D1- or A2A-Cre mice underwent stereotaxic surgery for infusion of a dilute Fox et al 2022 Cre-dependent eYFP virus in the NAc to label D1- or D2 MSNs, respectively. Tissue slices containing the NAc were collected after homecage fentanyl abstinence. (**H**) Representative image of sparsely labeled MSNs in the NAc, a single MSN after 3D reconstruction, and concentric rings for Sholl analysis. (**I**) Representative D1-MSNs and Sholl analysis from water and fentanyl abstinent mice (mean ± sem;*,p<0.05 Sidak’s post-hoc, n=48 cells from 18 mice). (**J**) Number of branch points and total dendritic length in D1-MSNs from water and fentanyl abstinent mice (**, p<0.005 nested t-test; n=48 cells from 18 mice; lines indicate median). (**K**) Representative D2-MSNs and Sholl analysis (mean ± sem), (**L**) number of branch points and total dendritic length in D2-MSNs from water and fentanyl abstinent mice. (p>0.05, n=47 cells from 17 mice; lines indicate median. See also Fig S1. Detailed statistics in Supplemental File 1.

We next asked how fentanyl abstinence altered dendritic morphology of medium spiny neuron (MSN) subtypes in the nucleus accumbens core (NAc). We used a low-titer Cre-dependent eYFP virus to sparsely label MSN subtypes in D1- and A2A-Cre mice (Fox et al., 2020a) (**Fig 1G-H;** fentanyl drinking data in **Fig S1B**). We found fentanyl abstinence decreased dendritic complexity of D1-MSNs as measured by reduced Sholl intersections (**Fig 1I**, p<0.05, at 40-70 µm from soma, Sidak’s post hoc after 2-way ANOVA), branch points (**Fig 1J**, p=0.005, nested t-test), and total dendritic lengths (**Fig 1J**, p=0.003, nested t-test). Dendritic complexity of D2-MSNs was unaltered by fentanyl abstinence as measured by equivalent Sholl intersections (**Fig 1K**, p>0.05), dendritic lengths (**Fig 1L**, p>0.05), and branch points (**Fig 1L**, p>0.05). There were no robust effects on dendritic spine density in either MSN subtype (**Fig S1C**, p’s>0.05).

To ensure changes to MSN morphology resulted from fentanyl abstinence and not fentanyl exposure alone, we performed similar experiments in eYFP-expressing D1- and A2A-Cre mice that did not undergo abstinence. These mice were perfused at a timepoint where bottles would normally be replaced with plain water. When we examined dendritic complexity of D1- and D2- MSNs in fentanyl exposed, but not abstinent mice, we found no significant changes to D1-MSN dendritic complexity compared to water exposed mice (**Fig S1D;** water vs fentanyl-no-abstinence, p’s >0.05 at all Sholl radiuses, number of branch points, total dendritic length). D2-MSN dendritic complexity remained unchanged by fentanyl (**Fig S1E**, p’s >0.05).

### Homecage fentanyl abstinence increases D1-MSN excitability and sEPSC amplitude

Reduced dendritic complexity is associated with altered excitability of D1-MSNs (Francis et al., 2017, 2019). Thus, we next assessed how fentanyl abstinence altered MSN physiology. We used AAV-DO-tdTomato-DIO-eGFP (Saunders et al., 2012) in D1-Cre mice to label Cre-positive D1-MSNs in green, and Cre-negative D2-MSNs in red (Timeline in **Fig 2A**, representative recording site in **Fig 2B**, drinking data for mice available in **FigS5H**). Since opioid-induced electrophysiologic adaptations are NAc subregion specific (Madayag et al., 2019), we restricted our analysis to cells in the NAc core, and at the core/shell boundary. Fentanyl abstinence did not alter the frequency of spontaneous excitatory post-synaptic currents (sEPSCs) onto D1-MSNs (**Fig 2C**, p=0.187, nested t-test). Instead, we found abstinence increased the median sEPSC amplitude relative to D1-MSNs from water mice (**Fig 2D** Left: p=0.011, nested t-test), and caused a rightward shift in the cumulative probability plot (**Fig 2D** Right: Kolmogorov-Smirnov D=0.14, p=0.01). Fentanyl abstinence did not alter median sEPSC inter-event-interval (**Fig 2E**, p=0.642, nested t-test), nor shift the cumulative probability (Kolmogorov-Smirnov D=0.01, p>0.99). Along with increased sEPSC amplitude, fentanyl abstinence also caused a hyper-excitable phenotype in D1-MSNs marked by a more hyperpolarized action potential threshold (**Fig 2F**, p=0.006, nested t-test) and a trend towards an increase in the number of current-evoked spikes (**Fig 2G** Left: drug p=0.09, mixed effects analysis; Right: p=0.07 nested t-test). Fentanyl abstinence did not alter sEPSC frequency onto D2-MSNs (**Fig 2H**, p=0.467, nested t-test) and did not significantly increase median sEPSC amplitude (**Fig 2I** Left: p=0.252, nested t-test), or cumulative probability (**Fig 2I** Right: Kolmogrov-Smirnov D=0.02, p>0.99). We found no differences in median sEPSC inter-event-interval in D2-MSNs (**Fig 2J** Left: p=0.624, nested t-test), nor shifts in the cumulative probability plot for D2-MSNs from fentanyl abstinent mice (**Fig 2J** Right: Kolmogorov-Smirnov D=0.098, p>0.99). D2-MSN excitability was unchanged by fentanyl abstinence as there were no differences in threshold potential (**Fig 2K**, p=0.14, nested t-test) or current evoked spikes (**Fig 2L** Left: drug p=0.299, RMANOVA; Right: p=0.494, nested t-test). In both MSN subtypes, fentanyl abstinence did not significantly change resting membrane properties (**Fig S2**).

**Figure 2.**
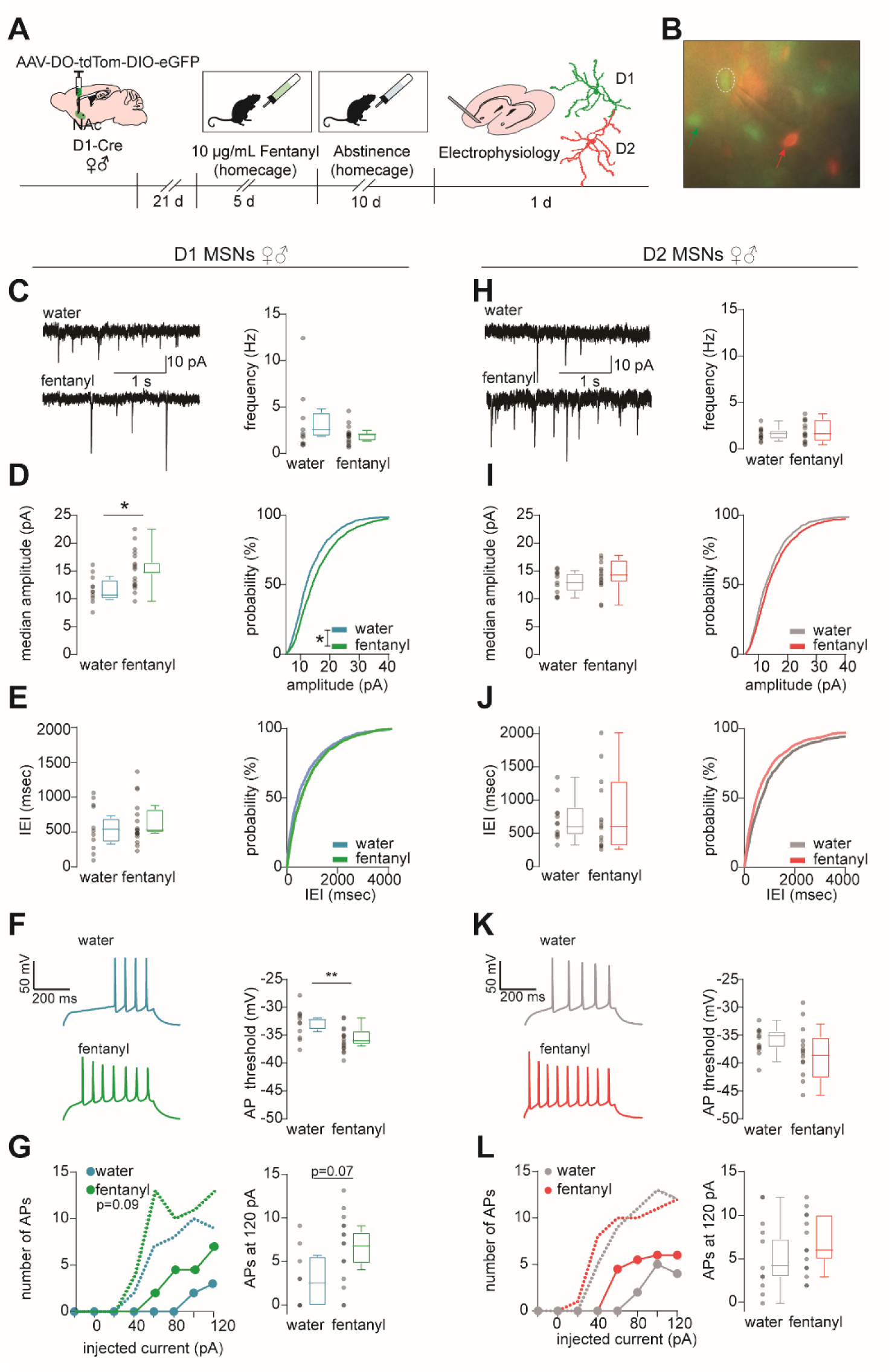
Fentanyl-abstinence increases D1-MSN excitatory input and excitability. **(A)** Schematic of MSN subtype labeling approach for electrophysiology experiments. Male and female D1-Cre mice received intra-NAc infusion of AAV-DO-TdTomato-DIO-eGFP virus so that Cre negative (i.e. D2-MSNs) express TdTomato, and Cre positive (D1-MSNs) express eGFP. Following homecage fentanyl abstinence, slices containing the NAc were collected for patch-clamp recording. (**B**) Representative image of MSN identification by red and green fluorescence prior to patching. Green and red arrows indicate D1- and D2-MSNs, respectively, patched D1-MSN denoted by hashed circle. (**C**) Spontaneous excitatory postsynaptic currents (sEPSCs) were recorded while holding the membrane potential at -50 mV and analyzed by template-based event detection. Representative sEPSCs and median sEPSC frequency, (**D**) median sEPSC amplitude and cumulative probability (*, p=0.012 nested t-test; *, p=0.01, Kolmogrov-Smirnov D=0.14), (**E**) median sEPSC inter-event-interval and cumulative probability in D1-MSNs from water and fentanyl abstinent mice. Individual data points represent individual neurons. Box and whiskers plots show the median and range for data nested by mouse (n=28 cells from 12 mice). (**F**) Representative evoked action potentials (APs) and the potential required to elicit an AP, (**G**) number of APs elicited by injecting current in 20 pA intervals (median + range, circles and dashed lines, respectively), and number of APs at 120pA in D1-MSNs from water or fentanyl abstinent mice (n=23 cells from 10 mice, *p*=0.07, nested t-test). (**H**) Representative sEPSCs and median sEPSC frequency, (**I**) median sEPSC amplitude and cumulative probability, (**J**) median sEPSC inter-event-interval and cumulative probability in D2-MSNs from water and fentanyl abstinent mice (n=26 cells from 18 mice). (**K**) Representative evoked APs and the membrane potential required to elicit an AP, (**L**) number of APs elicited by injecting current in 20 pA intervals (median + range), and number of APs at 120pA in D2-MSNs from water or fentanyl abstinent mice (n=25 cells from 17 mice). See also Fig S2. Detailed statistics in Supplemental File 1.

### Fentanyl abstinence causes distinct changes to D1- and D2-MSN translatomes

To determine the molecular mechanisms underlying D1-MSN specific atrophy and increased excitability, we extracted RNA from immunoprecipitated polyribosomes in D1- and D2 MSNs as in our previous work (Engeln et al., 2020; Fox et al., 2020a). We performed RNA sequencing of D1- and D2-MSN translatomes after fentanyl abstinence (**Fig 3A-B**, fentanyl drinking data in **Fig S3A**), and found very little overlap in the differentially expressed genes between MSN subtypes after fentanyl abstinence (**Fig 3C**). We used Nanostring to measure mRNA levels in D1- and D2-MSN translatomes and confirmed MSN subtype specificity of our samples (**Fig S3B**). Next, we performed Weighted Gene Co-Expression Network Analysis (WGCNA) in each MSN subtype. We identified 19 modules of co-expressed genes in D1-MSNs and 16 modules in D2-MSNs (**Fig 3D)**. From these 35 total modules, we chose 11 for further exploration based on p<0.05 for effect of fentanyl (**Fig 3E; Table S1**). The effects of fentanyl abstinence on the expression of these modules were primarily MSN-subtype-specific, except for the Black and Red modules which contained upregulated or downregulated genes, respectively, in both MSN subtypes. We next used Nanostring to assess differential expression of 3 randomly selected hub genes from each fentanyl module, and analyzed expression with sexes combined, as well as in each sex independently (**Fig 3G**). Despite reduced sensitivity for detecting small expression changes, our Nanostring analysis showed 16 of the 33 hub genes we selected were significantly differentially expressed in either one or both sexes, and all tested hub genes were concordant with directionality in the RNAseq data (i.e. up- or down-regulated by fentanyl abstinence). To further refine the fentanyl modules, we performed Gene Ontology (GO) Analysis on the genes from each module using BiNGO (Maere et al., 2005) similar to our previous work (Engeln et al., 2020) (**Supplemental File 2**). Given the changes to dendritic complexity and excitability, we looked for overrepresentation of synaptic and dendritic complexity GO terms (**Fig 4A**), and found enrichment in both Green and Magenta modules. We selected the Green module for further investigation since our dendritic complexity loss was specific to D1-MSNs, and the Green module contained genes downregulated in D1-MSNs (Green module structure in **Fig 3F**). We measured mRNA expression of additional genes in the Green module with Nanostring (**Fig 4B**), comparing both across and between the sexes. Several of the tested genes were significantly downregulated after fentanyl abstinence and were primarily restricted to D1-MSNs. Two of the tested Green module genes (Cdk5r1 and Nat8l) were significantly changed in D2-MSNs from female mice. When sexes were combined, we found significant downregulation of Gramd1b, Nat8l, and Prkce; when sexes were separated, downregulated Gramd1b and Prkce reached statistical significance in male mice, and Nat8l in female mice (**Fig 4B**).

**Figure 3.**
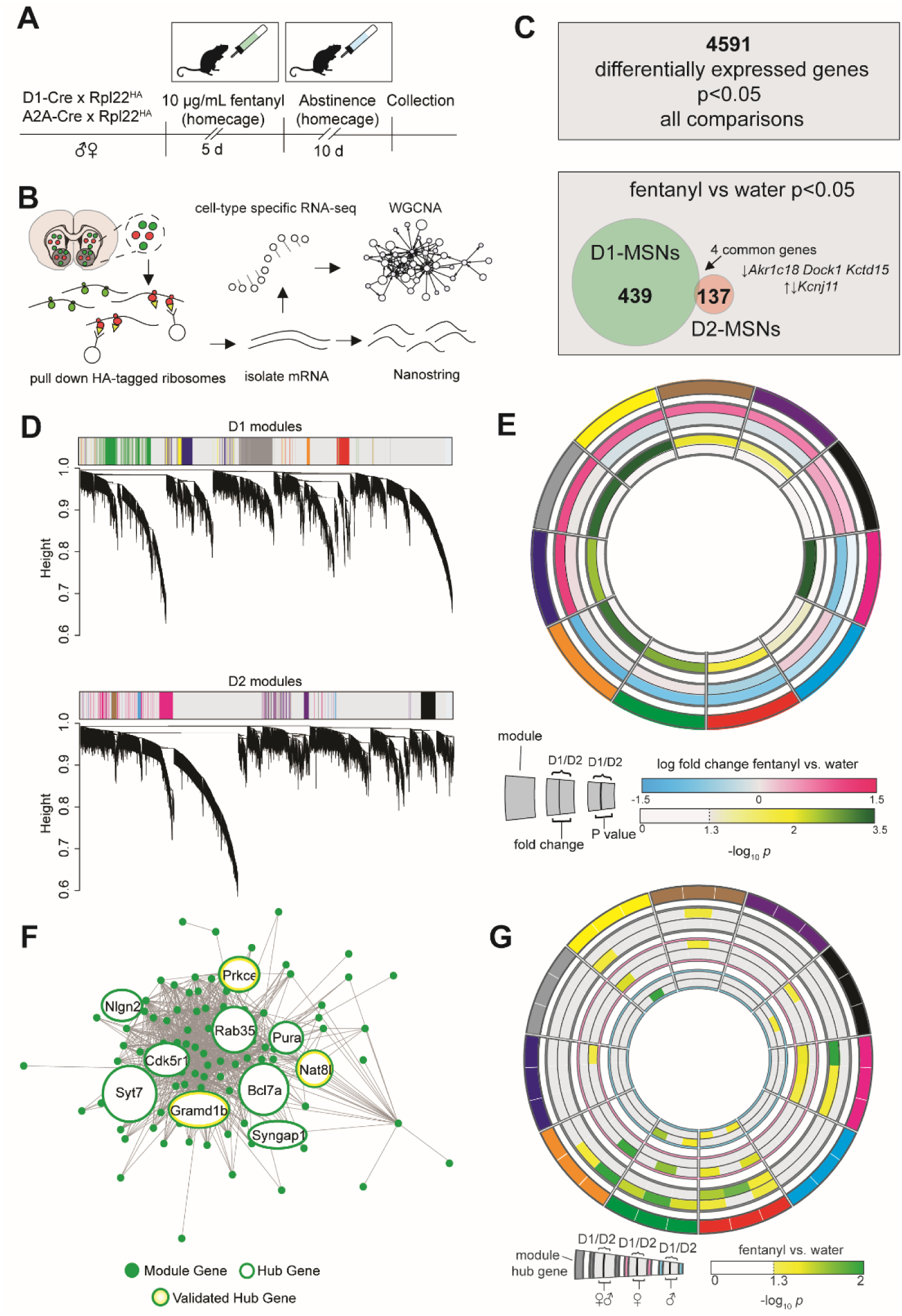
Translatome sequencing reveals MSN Subtype Specific Gene Co-expression networks in Fentanyl Abstinence. **(A)** Timeline for profiling MSN subtype translatomes in fentanyl abstinence. Male and female D1- and A2A-Cre mice were crossed with Cre-dependent ribosome tagged mice (Rpl22HA; RiboTag), then underwent homecage fentanyl drinking and abstinence (n=6 samples/sex/cell-type/drug, 4 mice pooled per sample). (**B**) HA-tagged ribosomes from D1- or D2-MSNs were immunoprecipitated and purified mRNA was used for Nanostring or to prepare cDNA libraries for RNAseq. Cell-type specific RNAseq data were analyzed by Weighted Gene Co-expression Network Analysis (WGCNA). (**C**) Top: number of genes that exhibited a nominally significant effect (p<0.05) of fentanyl, sex, or cell type. Bottom: number of genes with nominally significant effect of fentanyl in D1- and D2-MSNs showing little overlap between subtypes. (**D**) Clustering dendrograms from analysis of the 4591 genes in D1- and D2-MSNs with WGCNA. Module colors are shown for modules pertaining to fentanyl abstinence, defined as differential module eigengene expression with p<0.05 effect of fentanyl abstinence. (**E**) Circos plot showing 11 selected WGCNA modules as arbitrary colors (outermost ring), ranked clockwise by overall fold change in D1-MSNs. Fold change in eigengene expression, and –log_10_(*P* value) in D1- and D2-MSNs are shown internally. (**F**) Network structure of the green module. Hub genes are encircled in green, and the 3 hub genes selected for Nanostring are also encircled in yellow. Hub gene circle sizes are scaled based on the number of connections (**G**) Circos plot showing Nanostring analysis of 3 hub genes for each WGCNA module (outermost segments). Inner rings show fentanyl vs water differential expression scaled by –log_10_(*P* value) with significant (*P*<0.05) differences featured in color. The outer comparison rings show data with males and females combined, while the pink and blue encircled rings show comparisons in females or males only. See also Fig S3, Table S1. Detailed statistics in Supplemental File 1.

**Figure 4.**
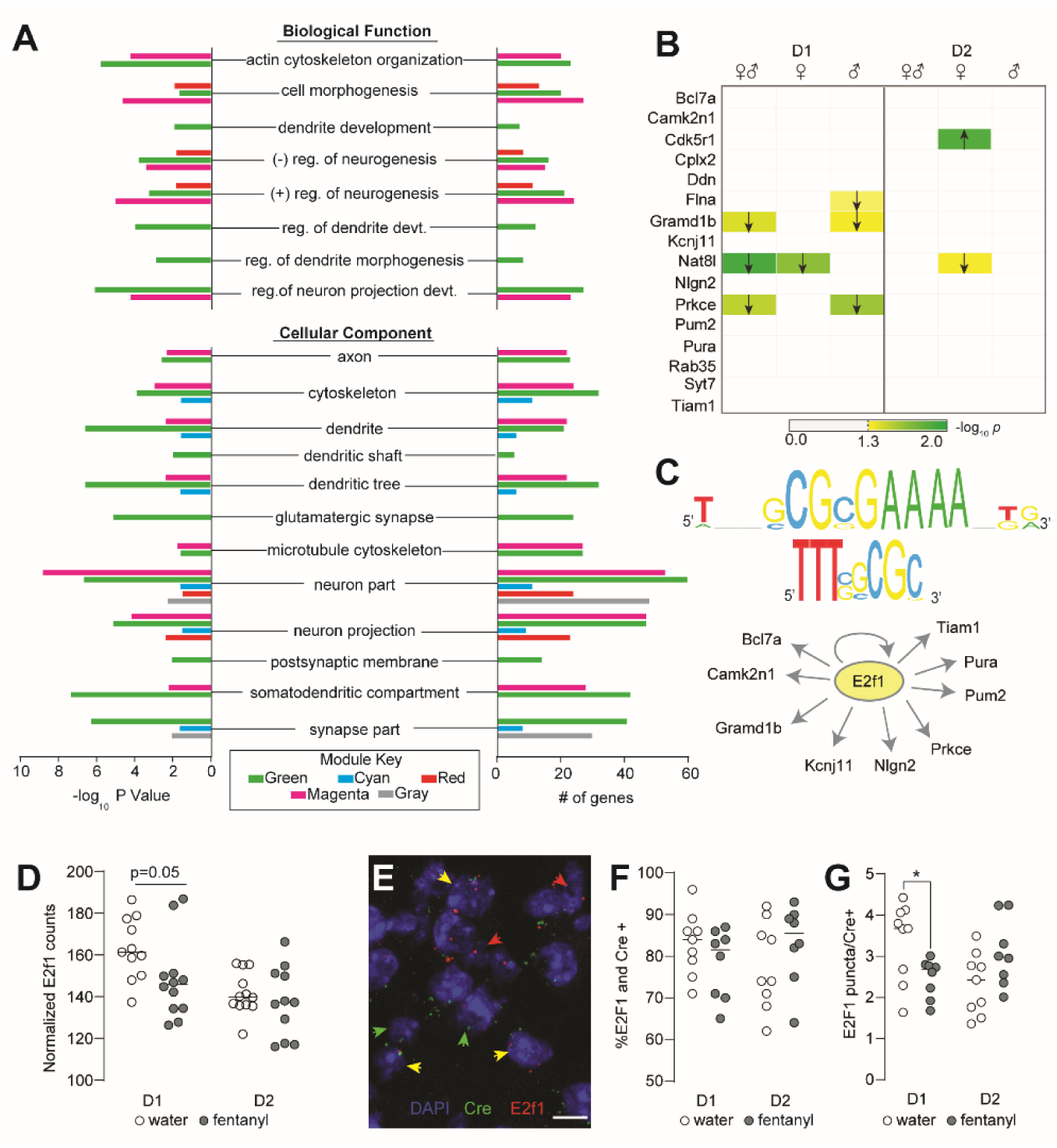
The D1-MSN specific Green module genes control dendritic morphology, are downregulated in D1-MSNs, and co-regulated by E2f1. **(A)** Gene Ontology analysis showing enrichment of synaptic and dendritic morphology related terms in green, cyan, red, magenta, and gray modules. (**B**) Nanostring validation of key green module genes in D1 and D2-MSNs showing differences in expression with sexes combined, and in males and females separately. Arrows denote direction of significant differences in expression between fentanyl and water. –log_10_(*P* value) is encoded in color. (**C**) Upstream regulator analysis with iRegulon indicates E2f1 is a predicted regulator of key green module genes. Predicted binding motifs transfac_public-M00024 and yetfasco 627 shown above. (**D**) Nanostring counts showing decreased expression of E2f1 in D1-MSNs of fentanyl mice (n=6 samples/sex/cell-type/drug, 4 mice pooled per sample, p=0.05, unpaired t-test). (**E**) Representative IMARIS reconstruction of fluorescence in situ hybridization for E2f1 in a D1-Cre mouse. DAPI stained nuclei are blue; red arrows denote E2f1 positive nuclei, green arrows denote Cre positive (D1 or D2), and yellow denote E2f1 and Cre positive nuclei. (Scale bar =10µm). (**F**) Percent E2f1 and Cre colocalized nuclei, and (**G**) number of E2f1 puncta per Cre positive nucleus in water and fentanyl abstinent mice. Each dot represents an individual mouse and is the sum of 3 separate images (∼600 Cre+ nuclei counted per mouse; *, *p*=0.028, unpaired t-test). Lines indicate median. See also Fig S3 and Supplemental File 2 and 4. Detailed statistics in Supplemental File 1.

### D1-MSN dendritic complexity molecules share common transcriptional regulators

Given the variation in the extent of Green module gene downregulation between the sexes, we next looked for potential upstream transcriptional regulators. We used iRegulon (Janky et al., 2014) to look for transcription factor binding motifs enriched in the promoters of Hub genes in the Green module. This analysis identified 13 enriched sequence motifs (**Supplemental File 4**). Of the transcription factors that recognize these motifs, E2F1 stood out because the *E2f1* gene was itself a member of the Green module, down-regulated following abstinence from fentanyl. Ten genes in the Green module were predicted as E2F1 target genes based on the evolutionarily-conserved enrichment of E2F1 sequence motifs +/-20 kb from their transcription start sites (**Fig 4C**; **Supplemental File 4**). From this gene list, we selected three predicted E2f1 targets for chromatin binding analysis with Cut & Run qPCR in mice that underwent fentanyl abstinence. Since this method used bulk NAc tissue, we did not detect any statistically significant changes, however there was a trend for increased E2f1 binding upstream of Nlgn2 in the fentanyl abstinent condition (**Fig S3E**, p=0.084, unpaired t-test).

Next, we sought to confirm our RNAseq results showing decreased E2f1 in D1-MSNs after analysis. Nanostring analysis revealed decreased E2f1 counts in fentanyl abstinence D1-MSN translatomes (**Fig 4D**, p=0.05), but not D2-MSN translatomes (p=0.49). We next replicated this finding with single molecule fluorescence in situ hybridization (RNAscope) in a separate cohort of D1- and A2A-Cre mice that underwent fentanyl abstinence (Drinking data available in **Fig S3C**). We used probes against E2f1 and Cre to identify E2f1 mRNA levels in NAc core D1-MSNs or D2-MSNs. Although fentanyl abstinence did not alter the number of E2f1 expressing Cre positive nuclei in either D1- or D2-MSNs, (**Fig 4F**, p’s >0.14) fentanyl abstinence was associated with a significant decrease in the number of E2f1 puncta per Cre-positive D1-MSN nucleus (**Fig 4G**, p=0.028). By contrast, there were no significant differences in D2-MSNs, and there was a trend towards increased E2f1 puncta in D2-MSNs from fentanyl abstinent A2A-Cre mice (p=0.068).

### Increased E2f1 in D1-MSNs protects against fentanyl abstinence-induced atrophy, plasticity, and behaviors

Since E2f1 was decreased in D1-MSNs from fentanyl abstinent mice, we made a Cre-inducible AAV to overexpress E2f1 in D1-MSNs during fentanyl exposure (virus validation in **Fig S4A-B**). In Cre-transfected Neuro2A cells, we found AAV-DIO-E2f1 decreased both mRNA and protein levels of E2f1 target genes (**Fig 5A**, mRNA: Nlgn2, p=0.047, Gramd1b p=0.0001; protein: Nlgn2, p=0.038, Gramd1b, p=0.044, unpaired t-tests). In D1-Cre mice expressing AAV-DIO-E2f1 (representative expression in **Fig 5B**), there were trends towards decreased expression of target genes Nlgn2 and Gramd1b in total NAc, likely due to E2f1 manipulation in only a subset of neurons (**Fig S4C**, p=0.085, p=0.09).

We next asked if E2f1 overexpression during fentanyl abstinence could protect against D1-MSN dendritic atrophy and altered physiology. We co-infused AAV-DIO-E2f1 and DIO-eYFP in D1-Cre mice three weeks before fentanyl exposure and abstinence (**Fig 5C**, drinking data in **Fig S4D**). We found no differences in dendritic morphology between D1-MSNs in water or fentanyl abstinent mice expressing DIO-E2f1/eYFP (Representative D1-MSNs in **Fig 5D**) as measured by Sholl intersections, number of branch points, and total dendritic length (**Fig 5E**, p’s>0.05). In fentanyl abstinent mice, D1-MSNs expressing eYFP had reduced dendritic complexity, but E2f1/eYFP D1-MSNs were indistinguishable from D1-MSNs from both groups of water mice (**Fig S4E**). We also found significantly more branch points in E2f1/eYFP expressing D1-MSNs vs eYFP D1-MSNs from fentanyl abstinent mice (**Fig S4E**, p=0.022, Sidak’s post-hoc). Together, these data indicate E2f1 prevented abstinence-induced dendritic atrophy in D1-MSNs. Next, we performed patch-clamp electrophysiology in D1-cre mice co-expressing AAV-DO-tdTomato-DIO-GFP and AAV-DIO-E2f1. In mice expressing E2f1/GFP, fentanyl abstinence no longer increased median sEPSC amplitude (**Fig S5C**, nested t-test p=0.49), nor threshold potential (**Fig S5E**, p=0.233) relative to water mice Together, these data indicate E2f1 overexpression reduces the magnitude of the electrophysiological changes caused by fentanyl abstinence.

Given that increased E2f1 expression protected against D1-MSN atrophy and changes to excitatory drive, we next asked if it would protect against abstinence-induced stress susceptibility and other negative-emotional behaviors. Male and female D1-Cre mice received intra-NAc AAV-DIO-E2f1 or AAV-DIO-eYFP three weeks before fentanyl drinking and abstinence (**Fig 5F**). A subset of male mice underwent a brief social defeat stress, while other male and female mice underwent EPM and three-chamber social interaction tests, each separated by 24 hr. We found increased E2f1 expression in D1-MSNs prevented stress-susceptibility in fentanyl abstinent mice subject to brief social stress (**Fig 5G**, eYFP: water vs fentanyl abstinence, p=0.003; E2f1: water vs fentanyl abstinence=0.493, Sidak’s after ANOVA). In mice tested in EPM and three-chamber social interaction, only fentanyl-eYFP mice showed increased anxiety-like behavior (**Fig 5H**, eYFP: water vs fentanyl-abstinence, p=0.002; E2f1: water vs fentanyl abstinence, p=0.88, Sidak’s post-hoc) and reduced social interaction (**Fig 5I**, eYFP: water vs fentanyl abstinence, p=0.008; E2f1: water vs fentanyl abstinence, p=0.98, Sidak’s post-hoc). Importantly, E2f1 did not block behavioral effects by significantly altering fentanyl consumption between mice receiving eYFP or E2f1 (**Fig S4D**).

**Figure 5.**
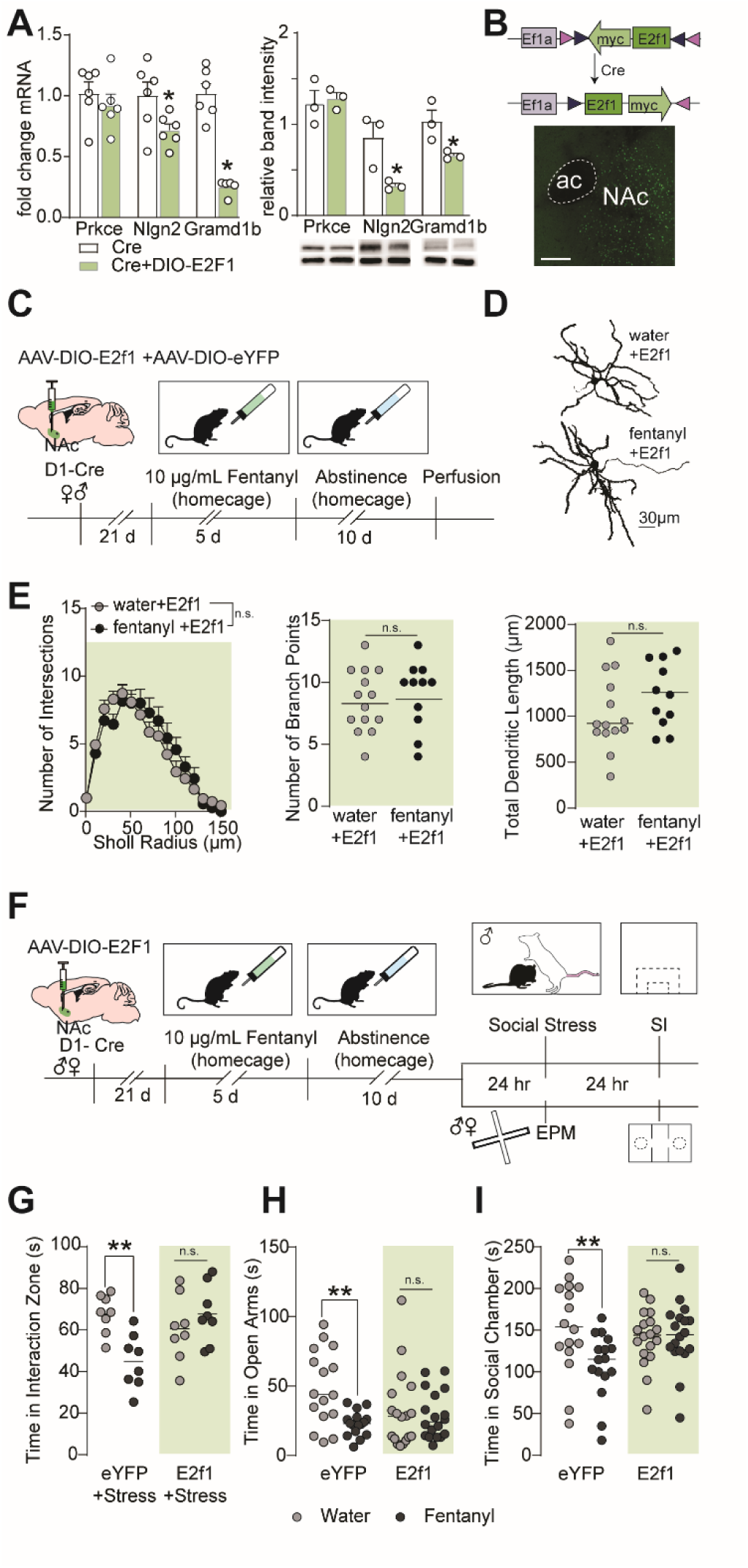
Increasing E2f1 in D1-MSNs protects against abstinence-induced atrophy and behavior. **(A)** Neuro2a cells were transfected with Cre and DIO-E2f1 or only Cre. Left: qPCR analysis showing downregulation of E2f1 target genes Nlgn2 and Gramd1b (*, p<0.05, n=6 cultures per condition). Right: western blot showing downregulation of Nlgn2 and Gramd1b at protein level. Top bands are the protein of interest and bottom are Gapdh used to normalize expression (n=6 cultures/condition pooled 2 cultures/sample. *, p<0.05). Bars are mean ± sem (**B**) Top: Cre-mediated translocation of E2f1-myc-flag vector allowing for D1-MSN specific expression. Bottom: Representative photomicrograph showing AAV expression in the NAc. (**C**) Timeline for E2f1 morphology experiments. Male and female D1-Cre mice received AAV-DIO-E2f1 and a dilute AAV-DIO-eYFP to label D1-MSNs after fentanyl abstinence (**D**) Representative D1-MSNs from mice receiving AAV-DIO-E2f1 prior to water or fentanyl abstinence. (**E**) Sholl analysis, number of branch points, and total dendritic lengths showing E2f1 prevents atrophy of D1-MSNs following fentanyl abstinence (Sholl: mean±sem, Sidak’s post-hoc after ANOVA, p>0.05; Nested t-tests, p’s>0.05, n=28 cells from 9 mice; lines indicated median). (**F**) Timeline for E2f1 behavioral experiments. Female and male D1-Cre mice received AAV-DIO-E2f1 or AAV-DIO-eYFP prior to undergoing fentanyl drinking and abstinence. A subset of male mice (top) was subjected to a one-day social stress paradigm, then assessed for stress-susceptibility 24hr later. The remaining mice underwent EPM and three-chamber social interaction testing. (**G**) Individual data points represent time stressed water and fentanyl abstinent mice spent in the interaction zone with a novel target present. (**, p=0.003, eYFP-water vs eYFP-fentanyl, Sidak’s post-hoc). (**H**) Individual data points showing time water and fentanyl abstinent mice spent in the open arms of the EPM and (**I**) in the social chamber with a social target present (**, p’s<0.008 eYFP-water vs eYFP-fentanyl, Sidak’s post hoc). Lines indicate median. See also Fig S4. Detailed statistics in Supplemental File 1.

## Discussion

Here we show fentanyl abstinence induces neuron subtype-specific structural, functional, and molecular changes in the NAc. First, we found fentanyl abstinence increases anxiety-like behavior, social avoidance, and stress-susceptibility that is associated with a loss of D1-MSN dendritic complexity. Second, fentanyl abstinence increased D1-, but not D2-MSN excitability, and increased excitatory input onto D1-MSNs. These structural and functional changes were associated with altered expression of unique gene networks in each MSN subtype, including downregulation of transcriptionally co-regulated dendritic complexity and synaptic genes in D1-MSNs. Finally, increasing expression of transcriptional regulator E2f1 in D1-MSNs attenuated structural, functional, and behavioral changes caused by fentanyl abstinence.

The main goal of this study was to determine how fentanyl abstinence impacts NAc medium spiny neurons. Since NAc MSNs are also important for the behavioral response to stress (Fox et al., 2020a), we provided fentanyl in the drinking water to minimize injection and handling stress. Here, and in our other work, we found mice will reliably consume fentanyl water in amounts similar to plain tap water (Franco et al., 2022). Because mice receive fentanyl as they consume water throughout the day, this method has the added benefit of mimicking intermittent drug use (vs continuous exposure) and minimizing repeated bouts of withdrawal between injections. We found fentanyl exposed mice exhibited increased naloxone precipitated withdrawal signs, suggesting our method produces a degree of opioid dependence. Similar to other studies on protracted opioid abstinence (Bai et al., 2014; Becker et al., 2017; Bravo et al., 2020), we also found decreased social behavior and increased anxiety-like behavior in both sexes of abstinent mice. Since drug exposure canalter HPA axis function (Wemm and Sinha, 2019), we also assessed stress-susceptibility in male mice, using a subthreshold social stressor sometimes termed “microdefeat”(Golden et al., 2011). This procedure measures stress-susceptibility in a short, open-field based social avoidance test, and in the absence of an additional manipulation, mice do not show social avoidance. If mice receive a “pro-stress” manipulation prior to subthreshold stress, they show increased social avoidance. Using this procedure, we found fentanyl abstinence with subthreshold stress, but not abstinence alone increased social avoidance and stress-susceptibility. The effect of fentanyl abstinence alone is less pronounced in the open-field based test because the arena, testing duration, and social target are different from the 3-chamber social interaction test. The decreased social interaction we see here in the 3-chamber test is concordant with our previous work showing reduced social interaction in this test after 56 hr of fentanyl abstinence (Franco et al., 2022).

Drugs with abuse potential alter NAc MSN morphology. Canonically, psychostimulants increase dendritic spine density, while opioids instead decrease dendritic spine density (Robinson and Kolb, 2004). Further, dopamine depletion has long been associated with a loss of MSN dendritic complexity and spine density (Ingham et al., 1989; Meredith et al., 1995), and opioid abstinence or withdrawal reduces NAc dopamine release (Acquas et al., 1991; Fox et al., 2017; Pothos et al., 1991; Rossetti et al., 1992). Opioid-induced spine loss and dendritic atrophy have been replicated across a number of morphine studies (Diana et al., 2006; Graziane et al., 2016; Leite-Morris et al., 2014; Matsubara et al., 1999; Robinson and Kolb, 2004; Spiga et al., 2005) (but see (Pal and Das, 2013); however comparatively fewer studies have examined dendritic morphology changes in specific MSN subtypes (Martin et al., 2019), especially following fentanyl. Here we found fentanyl abstinence reduced dendritic complexity in D1-MSNs, but not D2-MSNs. These findings mirror those of our work in chronic stress where only D1-MSNs show reduced dendritic arborization (Fox et al., 2020a; Francis et al., 2017). Unlike our stress work where D2-MSNs have increased dendritic spine density (Fox et al., 2020b), here we found spine density was relatively unchanged after fentanyl abstinence. It is possible that spine density changes occur at a different timepoint than what we assessed. Indeed, changes to spine density are time- and opioid-sensitive. Shortly after drug-induced reinstatement to heroin seeking, D1-MSN dendritic spine density is reduced (Martin et al., 2019). For morphine, density is increased in the NAc one day after acquiring conditioned place preference (Kobrin et al., 2016); unchanged after 5 days of non-contingent exposure, and decreased at 21 days withdrawal (Graziane et al., 2016). Further work is needed to establish the precise timeline for any changes to dendritic spines in MSN subtypes during fentanyl exposure and abstinence.

Opioid abstinence is also associated with physiological changes to NAc MSNs that depend on the duration. Acute (3-4 days) morphine abstinence is associated with decreased intrinsic excitability (Heng et al., 2008); in contrast, excitatory drive is potentiated after protracted (10-14 days) morphine abstinence (Wu et al., 2012, 2013). Several recent studies have assessed how morphine abstinence alters excitatory transmission in specific NAc MSN subtypes. Both 1- and 10-14 day morphine abstinence reduce excitatory input onto D2-MSNs in the NAc shell (Graziane et al., 2016; Hearing et al., 2016; Madayag et al., 2019; McDevitt et al., 2019). In D1-MSNs, 1-day morphine abstinence has little impact on excitatory input (McDevitt et al., 2019), while 10-14 days increases GluA2 lacking AMPA receptors and excitatory drive on D1-MSNs specifically in the shell (Hearing et al., 2016; Madayag et al., 2019). Unlike the work of Hearing and colleagues, here we found adaptations in NAc core D1-MSNs. There are several methodologic differences that may contribute to the discrepancy between our findings, including route of administration (repeated injections vs drinking) and opioid type (morphine vs fentanyl). We found 10-day fentanyl abstinence increases sEPSC amplitudes, hyperpolarizes the threshold potential, and produces a trend towards more spikes per current injection in D1-MSNs. Like altered dendritic morphology, the increased D1-MSN excitability in fentanyl abstinence is consistent with increased D1-MSN excitability following chronic psychosocial stress. However, stress and fentanyl abstinence do cause unique adaptations, as stress *decreases* excitatory input onto D1-MSNs (Francis and Lobo, 2017), while fentanyl abstinence increases the size of sEPSCs. Together our findings suggest plasticity that either increases D1-MSN excitability to compensate for a loss of excitatory inputs, or decreases dendritic complexity to compensate for overexcitation. Identifying which specific glutamatergic inputs are strengthened or weakened onto D1-MSNs (LeGates et al., 2018) may underlie the differences between stress and fentanyl abstinence, and requires further investigation. Surprisingly, we found no significant changes to D2-MSN excitability or excitatory drive. It is possible that changes to D2-MSN excitatory input are more robust at a different withdrawal timepoint as those seen in (Zhu et al., 2016).

Altered MSN morphology and physiology arise as a consequence of altered gene expression. Here, we found fentanyl abstinence altered both D1- and D2-MSN translatomes, but to a greater extent in D1-MSNs. Numerous studies have identified molecular changes in the NAc that arise during opioid abstinence, or as a consequence of opioid use (Cahill et al., 2018; Ferguson et al., 2013; Lefevre et al., 2020; Martin et al., 2019; Mayberry et al., 2022; Spijker et al., 2004; Sun et al., 2016; Townsend et al., 2021), including recent work in postmortem human brain (Seney et al., 2021). Some notable similarities arise between our findings, and those of recent publications: neuronal morphology related genes are downregulated across species and paradigm (Lefevre et al., 2020; Mayberry et al., 2022; Townsend et al., 2021). Like our work in abstinent mice, Mayberry et al found downregulated Green Module genes in the NAc of male morphine self-administering rats (e.g. Neurl4, Grik5, Shank3); Townsend and colleagues found decreased Green Module gene Camk2n1 in the NAc of female fentanyl self-administering rats. It is worth noting we also find downregulation of Nlgn2 in D1-MSNs, again, consistent with the response to chronic stress in mice (Heshmati et al., 2018). There are a number of differentially expressed genes in D2-MSNs, including genes in the D2-MSN specific Magenta Module. Since our dendritic atrophy phenotype was restricted to D1-MSNs, we chose to focus on gene expression changes in D1-MSNs, however the D2-MSN expression changes may reflect adaptations which protect against increased dendritic spine density on D2-MSNs as seen in stress (Fox et al., 2020b), or other dendritic complexity changes. There are several promising targets for future investigation in the Magenta Module, including 2 hub genes that are transcription factors (Arx, Dlx1). Recapitulating the changes to D2-MSN translatomes in D1-MSNs may also represent an additional opportunity to block synaptic changes and dendritic atrophy.

To establish a role for the Green Module in mediating D1-MSN dendritic atrophy, we selected transcription factor E2f1 for further investigation. Our iRegulon analysis predicted E2f1 regulated many Green Module genes (including 5 hub genes) and we found E2f1 was also downregulated by fentanyl abstinence. Further, globally disrupting E2f1-DNA binding increases anxiety-like behavior (Ting et al., 2014), consistent with our behavioral findings. The E2f family is also known to regulate gene expression following cocaine (Cates et al., 2018, 2019; Feng et al., 2014), and their transcriptional activity can be influenced by opioid receptor activation (Tencheva et al., 2005). Outside of the brain, E2f1 is primarily nuclear where it serves to regulate the cell cycle; however in post-mitotic neurons, E2f1 is found in the cytoplasm—especially in neuronal processes (Ting et al., 2014; Wang et al., 2010). Despite its odd localization, E2f1 does bind to several gene promoters in cortical neurons, most notably DNA repair related genes following DNA damage (Zhang et al., 2020). Here, we find increasing E2f1 expression in N2a cells results in decreased expression of two target genes Nlgn2 and Gramd1b. At first, this seems at odds with our finding that increased E2f1 in D1-MSNs protects against abstinence-induced dendritic atrophy, since it should theoretically work by *preventing* downregulation of those same genes. However, E2f1 function has been poorly characterized in the intact brain, and its function in neuronal cytoplasm is undetermined. In our iRegulon analysis, we also found E2f1 contains at least one E2f1 motif, and our viral expression may have induced self-regulation. Thus, E2f1 overexpression may protect against dendritic atrophy but potentially engages mechanisms of self-regulation to maintain homeostasis in D1-MSNs by dampening select E2f1 targets. Coincidentally, abstinence from morphine self-administration was recently shown to downregulate genes with E2f1 binding motifs (Mayberry et al., 2022), and may thus be a way to manipulate maladaptive gene expression for both natural and synthetic opioids.

Here we found preventing D1-MSN dendritic atrophy was protective against increased anxiety-like behavior, social avoidance, and stress-susceptibility generated by fentanyl abstinence. This is consistent with our prior work, showing D1-MSN atrophy mediates the behavioral response to chronic stress, and that preventing or reversing changes to dendritic morphology decrease stress-related behaviors (Fox et al., 2020a; Francis et al., 2017). We also found that in the absence of dendritic atrophy, the abstinence-induced changes to physiology were reduced. Increasing D1-MSN E2f1 expression normalized sEPSC amplitudes and decreased the extent of the excitability differences between water and fentanyl abstinent mice. These data agree with work in stress showing atrophy prevention also blocks changes to excitability and excitatory input (Francis et al., 2017). Our findings add to growing evidence for the importance of NAc D1-MSNs in the expression of affective behavior.Future work should build upon these discoveries to establish how D1-MSN changes influence opioid self-administration and relapse.

Taken together, our findings illustrate that even a single bout of protracted fentanyl abstinence can produce structural, functional, and molecular changes in the brain, and that these changes contribute to an increase in negative affective symptoms. Individuals with opioid use disorder oscillate between periods of drug use and abstinence, likely magnifying these neural adaptations. Targeting the molecular mechanisms during the transition from opioid use to abstinence may provide an important new therapeutic avenue. Intervention during this period may prevent some of the neurobiological changes, along with reducing negative affective symptoms that drive relapse and overdose.

## Supporting information

Key Resources Table

## Author Contributions

Conceptualization, MEF; Methodology, MEF, SAA, SMT, MKL; Formal Analysis, MEF, ABW, SAA; Investigation, MEF, ABW, DF, EC, CAC, ME, MDT, VMR; Resources, RC, SMT, MKL; Writing-Original Draft, MEF; Writing-Review & Editing, MEF, SMT, SAA, MKL; Visualization, MEF, ABW; Supervision, MEF, SMT, MKL; Funding Acquisition, MEF, SMT, MKL.

## Funding and acknowledgements

This study was funded by NIH K99/R00 DA050575 to MEF; R01MH106500, R01DA047943, R21DA052101, R21DA048554 and R01DA38613 to MKL; R01 MH086828 to SMT.

## Declaration of interests

The authors have no conflicts of interest to disclose

## STAR Methods

### RESOURCE AVAILABILITYz

#### Lead Contact

Further information and requests for resources and reagents should be directed to and will be fulfilled by Mary Kay Lobo

#### Materials Availability

The plasmid generated in this study will be made available upon reasonable request

#### Data and code availability

Data : RNA-seq data have been deposited at GEO and are publicly available as of the date of publication. Accession numbers are listed in the key resources table. Differential expression and WGCNA data have been deposited at Mendeley and are publicly available as of the date of publication. The DOI is listed in the key resources table. All other data reported in this paper will be shared upon request.

For review purposes: https://data.mendeley.com/datasets/snpmrt8fj3/draft?a=eb791487-c337-4305-ae00-61826519e141

Code:This paper does not report original code

### EXPERIMENTAL MODEL AND SUBJECT DETAILS

All experiments were performed in accordance with the Institutional Animal Care and Use Committee guidelines at the University of Maryland School of Medicine (UMSOM). Gonadally male and female mice were given food and water *ad libitum* and housed in UMSOM animal facilities on a 12:12 h light:dark cycle. Male and female C57Bl/6 mice bred at UMSOM were used for all experiments. Male CD-1 retired breeders (Charles River, >4 months) were used as aggressors for SDS. Male and female hemizygous D1-Cre (Line FK150) were used for morphology, electrophysiology, RNAscope, and E2f1 experiments. Male and female hemizygous A2A-Cre (Line KG139) were used for morphology and RNAscope experiments. Homozygous Rpl22^HA^ (“RiboTag”) (Sanz et al., 2009) mice expressing Cre-inducible Rpl22^HA^ were crossed to D1-Cre or A2A-Cre mouse lines to generate male and female D1-Cre X Rpl22^HA^ or A2A-Cre X Rpl22^HA^ mice and used for subtype-specific RNAseq and Nanostring. All transgenic mice were bred on C57Bl/6 background. Mice were 8-10 weeks of age during experiments and randomly assigned to groups.

#### Fentanyl abstinence

Male and female mice were pair housed across a perforated divider with a sex-matched mouse for the duration of the experiment (Franco et al., 2022). Each mouse was provided with a single 50 mL conical tube with a rubber stopper and ball-point sipper tube (Ancare) containing 10 µg/mL fentanyl citrate (Cayman #22659) in tap water as their sole liquid source for 5 days. Water control mice received plain tap water in identical 50 mL tubes. Fentanyl orwater consumption was determined by weighing the tubes daily, then normalized to individual mouse bodyweight. Following 5 days of fentanyl or water, the tubes were then replaced with bottles containing plain tap water for 10 days of abstinence. On Day 10 of abstinence, mice were either euthanized for morphology, electrophysiology, or molecular experiments, or transferred to individual housing for behavioral testing unless otherwise noted.

### METHOD DETAILS

#### Behavioral Testing

All video-tracking was conducted with TopScan Lite software (Cleversys, Reston, VA, USA). For 3-Chamber social interaction testing, mice were placed in an arena divided into 3 chambers by clear acrylic dividers. The two outer chambers contain wire mesh cups, while the central chamber is empty. The experimental mouse is placed in the central chamber of the arena with two empty cups and allowed to explore for 5 min. Then the experimental mouse is allowed to explore the arena for an additional 5 min with unfamiliar sex-matched adult conspecific in one of the wire mesh cups. The amount of time spent in the chamber containing the cups (empty or novel mouse) is measured with video tracking (Franco et al., 2022). For elevated plus maze, mice were placed in the center of the maze and their activity in open and closed arms was recorded over 5 min with video tracking (Fox et al., 2020a). For social stress (Fox et al., 2020a; Francis et al., 2017) male mice were placed in hamster cages with perforated plexiglass dividers containing a novel, aggressive male CD1-resident. We used only male mice in the social stress paradigm since unmanipulated CD-1s will attempt to mate with female mice. Mice were defeated for 3 min by 3 different residents on a single day, each session separated by 15 min of sensory interaction. 24 hr later, social avoidance was assessed with video-tracking. Experimental mice were placed in an open field containing a perforated box. Time spent around the box (“interaction zone”) was compared between two trials (2.5 min each) during which the box was empty or contained a novel CD-1. For naloxone precipitated withdrawal signs, mice that still had access to fentanyl were injected with 1mg/kg naloxone (Luster et al., 2020), then placed into an empty glass cylinder. A blinded experimenter scored withdrawal signs for 10 min and determined bodyweight loss after 1hr. Points were assigned as previously (Uddin et al., 2021). (Graded signs: wet dog shakes: 1-2, 2 points; 2+, 4 points; Escape jumps: 2-4, 1 point, 5-9, 2 points, 10+, 3 points; paw tremors: 1-2, 2 points, 3+, 4 points; bodyweight loss in 60 min, 1 point per each 1%; Checked Signs: diarrhea, swallowing movements, teeth chattering, all 2 points each; genital grooming, abnormal posture, 3 points each). All behavioral experiments were conducted during the light cycle.

#### Stereotaxic surgery

At 5-6 weeks of age, D1-Cre mice were anesthetized with isoflurane (4% induction, 1.5% maintenance) and affixed in a stereotaxic frame (Kopf Instruments). An incision was made in the scalp, and holes were drilled to target the NAc core (AP+1.6mm, ML: ± 1.5 mm, DV: -4.4 mm,10º) (Fox et al., 2020a). 300-400 nL of Cre-inducible, double inverted open (DIO) reading frame adeno-associated viruses (AAVs) were infused bilaterally with Neuro Syringes (Hamilton). The scalp was closed with Vet Bond (3M), and mice were group housed for 3 weeks to allow for recovery and expression.

#### Immunostatining

Mice were transcardially perfused with 0.1M PBS and 4% paraformaldehyde. Brains were removed and post-fixed for 24 hr, then sectioned to a thickness of 100 µm with a vibratome (Leica, Germany). Slices were washed 3 × 5 min with PBS and blocked for 30 min in in PBS with 3% normal donkey serum (NDS) and 0.3% Triton X-100. Slices were incubated at 4ºC overnight in chicken anti-GFP (1:500; Aves Lab, Tigard, OR, USA; #GFP-1020).

Slices were washed 3 × 5 min, then 7 × 60 min, then incubated at 4ºC overnight in Anti-Chicken Alexa 488 (1:500; Jackson Immuno, West Grove, PA, USA; #111-545-144). Slices were washed with PBS as above, mounted with Vectashield, and imaged on a laser-scanning confocal microscope (Leica SP8).

#### MSN reconstruction and dendrite analysis

D1-Cre or A2A-Cre mice were injected with AAV5-Ef1a-DIO-eYFP (UNC Vector Core, Chapel Hill, NC, USA, diluted to 1.5 ×10^11^ VP/mL). Sections containing NAc were sampled from bregma AP:1.5-1.0 mm and Z-stack images were acquired at 0.6 µm increments using a 40x objective. MSNs were 3D reconstructed using Imaris 8.3 software (Bitplane, Oxford Instruments) as previously (Engeln et al., 2020; Fox et al., 2020a). Surfaces were masked togenerate a 2D image of a single MSN for Sholl analysis. Concentric ring intersections were determined using the ImageJ Sholl analysis plugin (Ferreira et al., 2014) at 10 µm increments from soma.

#### Electrophysiology

For electrophysiological recordings, D1-Cre mice expressing AAV9-Ef1a-DO-TdTomato-DIO-EGFP (Saunders et al., 2012) were deeply anesthetized with isoflurane and perfused with ice cold NMDG-substituted artificial cerebrospinal fluid (ACSF) prior to decapitation. NMDG-ACSF contained (in mM) 92 N-methyl-D-glucamine, 2.5 KCl, 1.25 NaH_2_PO_4_, 30 NaHCO_3_, 20 HEPES, 25 glucose, 0.5 CaCl_2_ and 10 MgCl_2_. The brain was dissected out and 300 µm thick coronal slices containing nucleus accumbens were cut in ice-cold NMDG-ACSF with a vibratome. Slices were then placed in interface slice chambers at room temperature and allowed to rest for at least an hour before recording in Na-ACSF containing (in mM) 120 NaCl, 3 KCl, 1.0 NaH_2_PO_4_, 1.5 MgSO_4_·7H_2_O, 2.5 CaCl_2_, 25 NaHCO_3_, and 20 glucose. Pipettes with access resistance between 2-6 MΩ were filled with a K-gluconate based internal pipette solution containing (in mM) 105 K-gluconate, 5 KCl, 2 MgCl_2_·6H_2_O, 10 HEPES, 4 Mg-ATP, 0.3 Na-GTP, 1 EGTA, and 10 Na_2_-Phosphocreatine. Whole-cell recordings of D1 or D2 medium spiny neurons were done in the nucleus accumbens core. Cell type identification was done via fluorescent imaging while patching with D1- and D2-expressing MSNs expressing GFP and TdTomato respectively. Following patching, 5 minutes was allowed for the internal pipette solution to fill the cell before recordings were performed. Recordings were performed using CV203 BU headstage, Axopatch 200B amplifier, Digidata 1440A digitizer and Clampex 10.5 software (Axon Instruments). If the cell had an access resistance above 25 MΩ, it was excluded. Spontaneous excitatory postsynaptic currents (sEPSCs) were recorded in the absence of TTX while holding the membrane potential at -50 mV and analyzed by event detection in MiniAnalysis (Synaptosoft). All other electrophysiological data were analyzed in Clampfit 11.2. Intrinsic membrane properties were recorded by applying a 500 ms long -5 mV step from a holding potential of -70 mV. Access resistance was determined by the peak of the capacitive current deflection, membrane resistance by the mean current deflection after the cell had reached a steady state, and capacitance by the area of the current deflection. The resting membrane potential was recorded in current-clamp mode, while applying no holding current. Cell excitability was recorded by applying 500 ms long 20 pA current steps and counting the number of evoked action potentials. The threshold potential was measured as the membrane potential where the potential began to deflect to initiate the first recorded action potential. The Rheobase was recorded by applying a 300 pA current ramp and measuring the current required to evoked the first action potential. Current-clamp recordings were performed while using a holding current that kept the membrane potential at -70 mV at rest. We analyzed cells located only in the NAc core and the core/shell boundary. Cells were excluded from the experiment if they failed Grubb’s Outlier test (α=0.05) for a given intrinsic measurement.

#### RNA isolation

Polyribosomes were immunoprecipitated from NAc of D1-Cre-RiboTag and A2A-Cre-RiboTag mice as described previously (Chandra et al., 2017; Engeln et al., 2020; Fox et al., 2020a; Francis et al., 2017). Briefly, four 14-gauge NAc tissue punches per mouse (4 mice pooled/sample) were collected on Day 10 of fentanyl abstinence. Tissue punches were homogenized by douncing in 1 mL homogenization buffer, and the supernatant was incubated with 5 µl anti-HA antibody (Covance, Princeton, NJ, USA: #MMS-101 R) at 4ºC overnight with constant rotation. Samples were then incubated with 400 uL of protein G magnetic beads (Life Technologies, Carlsbad, CA, USA #100.09D) overnight at 4ºC with constant rotation. Beads were washed in a magnetic rack with high-salt buffer. RNA was extracted with a DNase step (Qiagen, Germantown, MD, USA) using the RNeasy mini kit (Qiagen #74104) by adding lysis buffer supplemented with beta mercaptoethanol and following the manufacturer’s instructions. RNA concentration and quality was determined with a Bioanalyzer (Agilent).

#### RNAseq and WGCNA

For RNA sequencing, only samples with RNA integrity numbers >8 were used. 6 samples/sex/drug/cell-type were submitted for RNA sequencing at the UMSOM Institute for Genome Sciences (IGS) and processed as described previously (Engeln et al., 2020). Libraries were prepared from 10 ng of RNA from each sample using the Smart-Seq v4 kit (Takara). Samples were sequenced on an Illumina HiSeq 4000 with a 75 bp paired-end read. An average of 64–100 million reads were obtained for each sample. Reads were aligned to the mouse genome (Mus_musculus.GRCm38) using TopHat (version 2.0.8; maximum number of mismatches = 2; segment length = 30; maximum multi-hits per read = 25; maximum intron length = 50,000). The number of reads that aligned to the predicted coding regions was determined using HTSeq. We applied two strategies to characterize gene expression changes in these data. First, we sought to identify individual genes with significant gene expression changes following abstinence from fentanyl. We used limma-trend to fit log2-normalized read counts per million to a linear model and tested for significant effects of fentanyl in each cell type, separately in males and in females, as well as treating sex as a covariate. Second, we characterized the effects of fentanyl on gene co-expression networks. As a starting point for this analysis, we used limma-trend to select the set of genes that exhibited a nominally-significant effect (p-value < 0.05) of fentanyl, sex, or cell type. We then used weighted gene co-expression network analysis (WGCNA) to characterize co-expressed modules among these genes, separately within D1 and D2 MSNs. Module detection was performed with the blockwiseModules() function with power = 10, corType = ‘bicor’, networkType = ‘signed’, minModuleSize = 25, reassignThreshold = 0, mergeCutHeight = 0.25, minMEtoStay = 0, and otherwise default parameters. We tested for differential expression of module eigengenes with limma, using posthoc contrasts to estimate effects of fentanyl while adjusting for sex differences, as above. Hub genes for each module were defined by calculating the Pearson correlation between the eigengene and each gene in the module.

#### Nanostring

40 ng of RNA per sample was processed with the nCounter Master Kit (Nanostring Technologies) by UMSOM IGS on a custom-made gene expression Code set. (Sequences available in **Supplemental File 4)**. Data were analyzed using cell-type, treatment, and sex as factors with nSolver Analysis software (Nanostring Technologies, RRID:SCR_021712) using Aars, Pum1, and Hsp90ab1 as housekeeping genes.

#### Gene Ontology and transcription factor analysis

We performed Gene Ontology (GO) Analysis using the BINGO plugin for Cytoscape (Maere et al., 2005) using the mouse annotation from geneontology.org in August 2019. We queried over-represented GO terms in each WGCNA module under the “molecular function,” and “cellular-component” ontologies using the whole annotation as a reference set, and a Benjamini & Hochberg False Discovery Rate (FDR) correction of 0.05. We identified common transcriptional regulators of the Green module using the iRegulon plug-in in Cytoscape (Janky et al., 2014) using program default settings with the exception of a more stringent enrichment score threshold of 3.5.

#### Cell-type specific E2f1 manipulation

Mouse E2f1-myc-Flag ORF clone was obtained from Origene (#MR206856). We PCR amplified the ORF using Phusion DNA polymerase (New England Biolabs #E0553), then cloned into the NheI and AscI restriction site of pAAV-Ef1a-DIO-eYFP (Addgene #27056). Vector sequences were confirmed by digestion and sequencing. AAV9-Ef1a-DIO-E2f1-myc-Flag was packaged at the University of Maryland Virus Vector Core Facility.

#### In Vitro Experiments

Neuro2A cell cultures were maintained in DMEM supplemented with 10% FBS and Penicillin/streptomycin (all from Thermo Fisher Scientific). Cell cultures of passage number between 5-10 were used for the entire experiment. 24 hours prior to transfection, cells were seeded at the density of 1.5 × 10^5^ cells per well in 24 well plates. Transfection was performed using Invitrogen LF3000 reagents following the manufacturer’s standard protocol. Cells were harvested 48 hours post transfection for either RNA or protein sample preparation.

For western blotting, cell lysates were prepared from 12 wells (2 pooled/condition for N=3 samples each) with RIPA buffer (Sigma-Aldrich R0278) supplemented with Phosphatase Inhibitor Cocktail 2 (Sigma-Aldrich P5726), Phosphatase Inhibitor Cocktail 3 (Sigma-Aldrich P0044), Complete Protease Inhibitor Cocktail (Roche 11873580001). Electrophoresis was performed using Mini-Protean TGX Gels (Biorad 4569033) followed by the transfer onto a nitrocellulose membrane (Biorad 1620094). After blocking in 5% skim milk/Tris-buffered saline supplemented with 0.1% Tween 20 (TBST), membranes were incubated with primary antibody overnight at 4°C followed by secondary antibody incubation for 1 hour at room temperature. Blot images were captured using Biorad ChemiDoc MP Imager, and the band intensities were quantified relative to GAPDH band intensities using ImageJ (NIH). The following primary antibodies were used at 1:1000 dilution: E2F1 (Abcam ab137415), PRKCE (Proteintech 20877-I-AP), GRAMD1B (Proteintech 24905-I-AP), NLGN2 (ProSci 7969), GAPDH (Cell Signaling 2118s).

For qPCR, RNA extraction samples were prepared from 12 wells (6 samples/condition) using the Qiagen RNeasy micro plus kit (Qiagen 74034). Neuro2a cells were harvested directly from the wells with application of Buffer RLT and physical scraping using a cell scraper. Purified RNA was quantified with a Nanodrop (Thermo), then 500ng of complementary DNA (cDNA) was synthesized using the reverse transcriptase iScript complementary DNA synthesis kit (Bio-Rad, Hercules, CA; # 1708891), then diluted to a concentration of 2.5 ng/µL. Relative mRNA expression changes were measured by quantitative PCR using Perfecta SYBR Green FastMix (Quantabio, Beverly, MA; #95072) with a Bio-Rad CFX384 qPCR system. Primer sets are available in the Key Resources **Table**. Fold change expression was calculated using the 2^-ddCt^ method with *Gapdh* as the reference gene.

#### Cut & Run qPCR

Nucleus Accumbens tissue punches were collected on abstinence Day 10, flash frozen on dry ice, and stored at -80°C. Cut & Run was performed using Cell Signaling Cut & Run Kit (#86652) following the manufacturer’s standard protocol. Tissue punches were homogenized by douncing in buffer supplemented with spermidine and protease inhibitor cocktail according to the manufacturer’s recipe. Enriched chromatins were purified using Cell Signaling ChIP and Cut & Run DNA Purification Kit (14209) following the manufacturer’s standard protocol. The following antibodies were used at 2ul per sample for negative control and 5ul per sample for all remaining antibodies: Normal Rabbit IgG (Cell Signaling #66362, negative control), H3K4me3 (Cell Signaling #9751, positive control), E2F1 (Invitrogen #32-1400). Expression changes were measured using qRT-PCR with PerfectaSYBR Green FastMix (Quanta, Beverly, MA, USA). Samples were normalized to negative control and compared with IgG. Samples were excluded if the positive control did not show enrichment relative to negative control. Primer sequences can be found in the Key Resources **Table**.

### QUANTIFICATION AND STATISTICAL ANALYSIS

Mice with off-target virus injection sites were excluded from analysis. All statistical tests were performed in Graph Pad Prism 9 and JASP (https://jasp-stats.org/) accounting for sex as a biological variable with ANOVA. Because there were a limited number of significant sex effects or sex interactions (α=0.05) the sexes were combined except where noted. To account for multiple cells from the same animal, we used nested t-tests. We used mixed-effects analysis to analyze the number of action potentials at a given current injection instead of RM-ANOVA due to one missing value. For nanostring and RNAscope differences, we used unpaired t-test independently in each cell type. A detailed table containing all statistical analysis, including sex effects and any normality corrections is in **Supplemental File 1**. Sample sizes were determined based on prior literature.

**Supplemental File 1**. Detailed statistical analysis including accounting for sex as a biological variable. Associated figure, comparison groups, and statistical tests used. N’s represent number of individual mice, n’s represent number of individual cells. The F, T, or K-S D column contains the given source of variation used in ANOVA, or specifies t for t-tests and K-S for Kolmogorov Smirnov Distance. P values and corresponding significance stars are shown for each comparison. Notes specify instances in which only certain comparisons are shown for brevity’s sake.

**Supplemental File 2**. Related to Figure 4. Enriched Gene Ontology terms for 11 selected WGCNA modules

**Supplemental File 3**. Nanostring CodeSet information

**Supplemental File 4**. Related to Figure 4. Transcriptional regulation analysis, predicted E2f1 target genes and motif ID’s from iRegulon.

## Notes

### Competing Interest Statement

The authors have declared no competing interest.

