## Supplemental Figures for "Negative emotional behavior during fentanyl abstinence is mediated by adaptations in nucleus accumbens neuron subtypes"

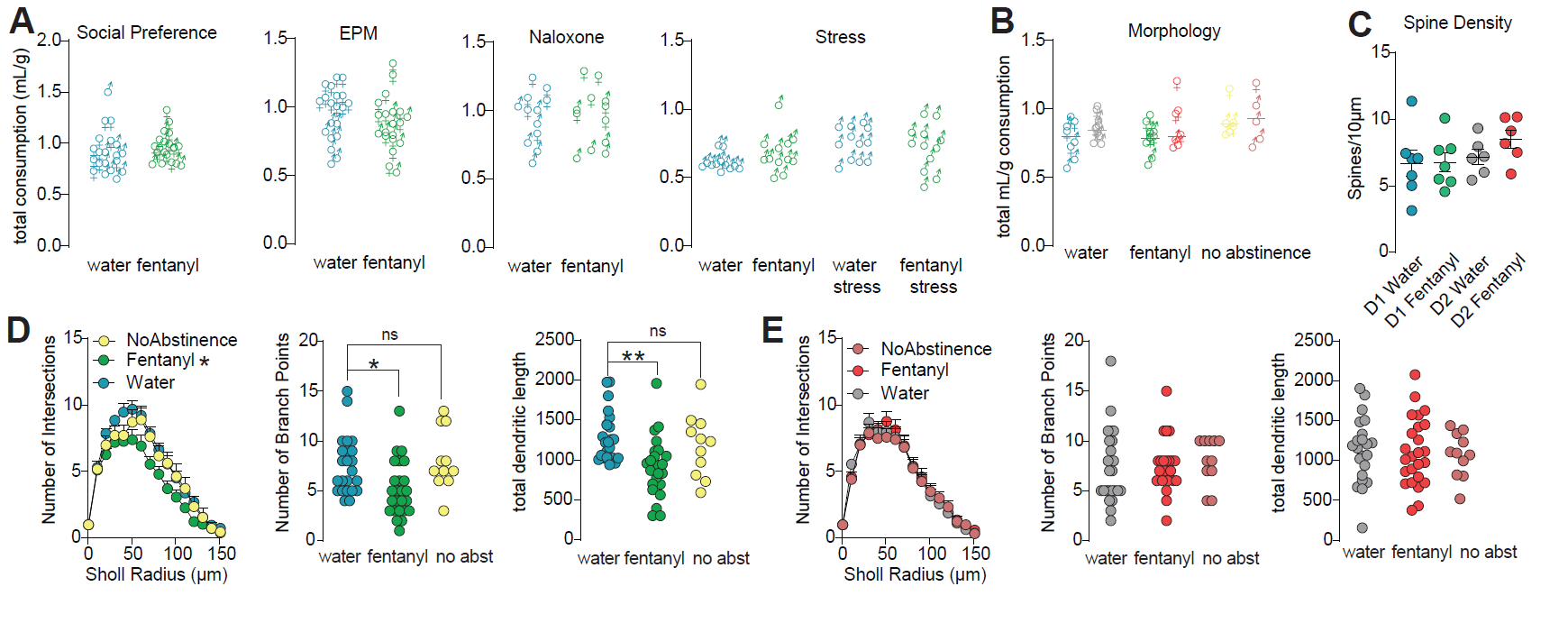

**Figure S1**. **Related to Figure 1.**

(**A**) Total liquid consumption across 5 days of water or fentanyl normalized to individual mouse bodyweight in male (♂) and female (♀) C57BL/6 mice used in behavioral experiments reported in Figure 1. (**B**) Total liquid consumption across 5 days of water or fentanyl normalized to individual mouse bodyweight in male and female D1- and A2A-Cre mice used in morphology experiments reported in Figure 1. (**C**) Spine density from a subset of water and fentanyl abstinent D1- and D2-MSNs. (**D**) Comparison of D1-MSN dendritic complexity between D1-Cre mice exposed to 5 days of fentanyl (no abstinence), and the water or fentanyl abstinent D1-Cre mice from Fig 1. (Dunnet’s post-hoc: No abstinence vs water p’s>0.05 for all measures, Fentanyl abstinence vs water p’s<0.05, n=59 cells from 24 total mice). (**E**) Comparison of D2-MSN dendritic complexity between A2A-Cre mice exposed to 5 days of fentanyl (no abstinence), and the water or fentanyl abstinent A2A-Cre mice from Fig 1 (All p’s >0.05, n=58 cells from 22 total mice). Detailed Statistics in Supplemental File 1.

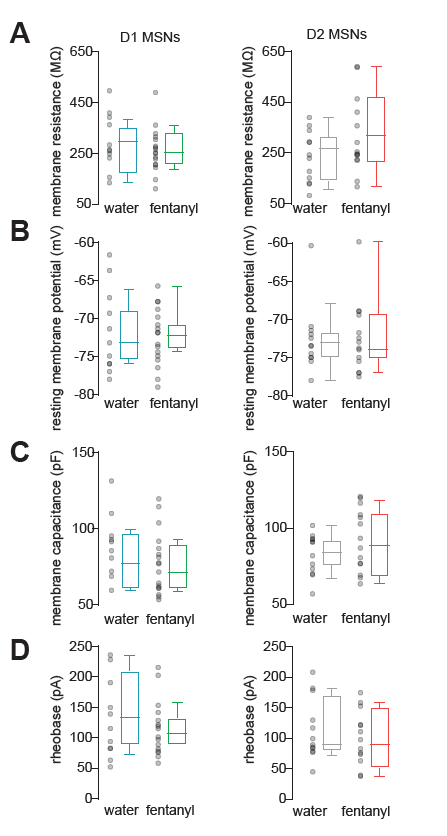

**Figure S2. Related to Figure 2**

**(A)** Membrane resistance, (**B**) resting membrane potential, (**C**) membrane capacitance, and (**D**) rheobase in D1- and D2-MSNs from water and fentanyl abstinent mice. Individual data points represent individual neurons. Box and whiskers plots show the median and range for data nested by mouse. Detailed Statistics in Supplemental File 1.

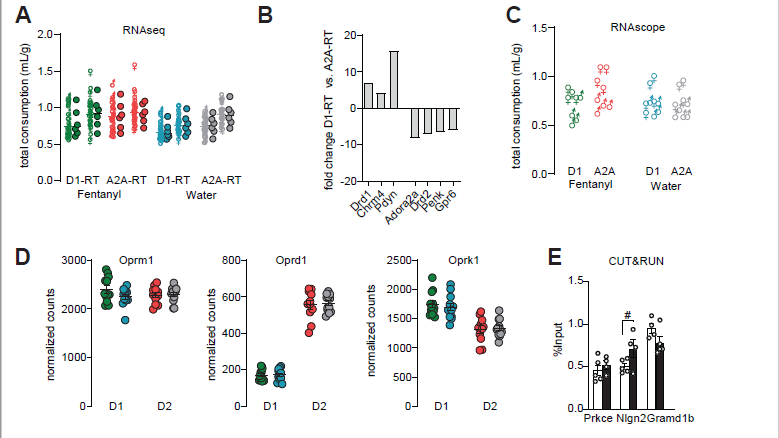

Figure S3. Related to Figures 3 and 4.

**(A)** Total liquid consumption across 5 days of water or fentanyl normalized to individual mouse bodyweight in male (♂) and female (♀) D1-RiboTag (RT) and A2A-RT mice used in RNAseq and Nanostring experiments in Figures 3 and 4. Tissue from mice that consumed similar amounts were pooled to generate 6 samples/sex/condition/genotype, resulting in an average consumption value per sample (filled circles next to individual points). (**B**) Nanostring validation showing expression of D1-MSN specific genes Drd1, Chrm4, and Pdyn is enriched, and D2-MSN genes Adora2a, Drd2, Penk, and Gpr6 expression is impoverished in D1-RT samples. (**C**) Total liquid consumption across 5 days of water or fentanyl normalized to individual mouse bodyweight in male (♂) and female (♀) D1 and A2A-Cre mice used for RNAscope experiments in Figure 4. (**D**) Nanostring counts of mu, delta, and kappa opioid receptors in samples from water and fentanyl abstinent D1- and A2A-RTmice. (**E**) Cut&Run chromatin profiling in total NAc from individual water (white) and fentanyl abstinent (black) mice. Regions containing predicted E2f1 binding sites for Prkce, Nlgn2, and Gramd1b were amplified with qPCR. Data are presented as (mean ± sem) expression in E2f1-immunoprecipitated samples relative to input. (#, p=0.08 water vs fentanyl abstinence for Nlgn2). Bars are mean ± sem. Detailed Statistics in Supplemental File 1.

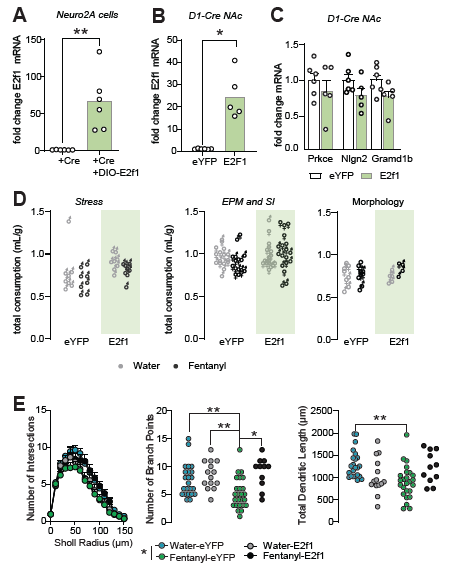

**Figure S4. Related to Figure 5.**

(**A**) Fold E2f1 expression in Neuro2A cells transfected with Cre only, or Cre and DIO-E2f1. (**, p=0.010, t-test). (**B**) Fold E2f1 expression in total NAc of D1-Cre mice expressing DIO-eYFP or DIO-E2f1. (**, p= 0.007, t-test). (**C**) Fold expression of E2f1 target genes in total NAc of D1-Cre mice expressing DIO-eYFP or DIO-E2f1. (Prkce, p=0.4, Nlgn2, p= 0.1, Gramd1b, p=0.08, t-tests). (**D**) Total liquid consumption across 5 days of water or fentanyl normalized to individual mouse bodyweight in male (♂) and female (♀) D1-Cre mice used in behavior and morphology experiments in Fig 5. (**E**) Comparison of D1-MSN dendritic complexity between fentanyl and water abstinent D1-Cre mice with Intra-NAc DIO-E2f1 or DIO-eYFP. (Sholl: *, p=0.014, Fentanyl-eYFP vs Water-eYFP; Branch points: **, p<0.002, *, p=0.022; Dendritic length: **, p=0.002; All Sidak’s post-hoc after ANOVA). Bars and Sholl analysis are mean ± sem. Detailed Statistics in Supplemental File 1.

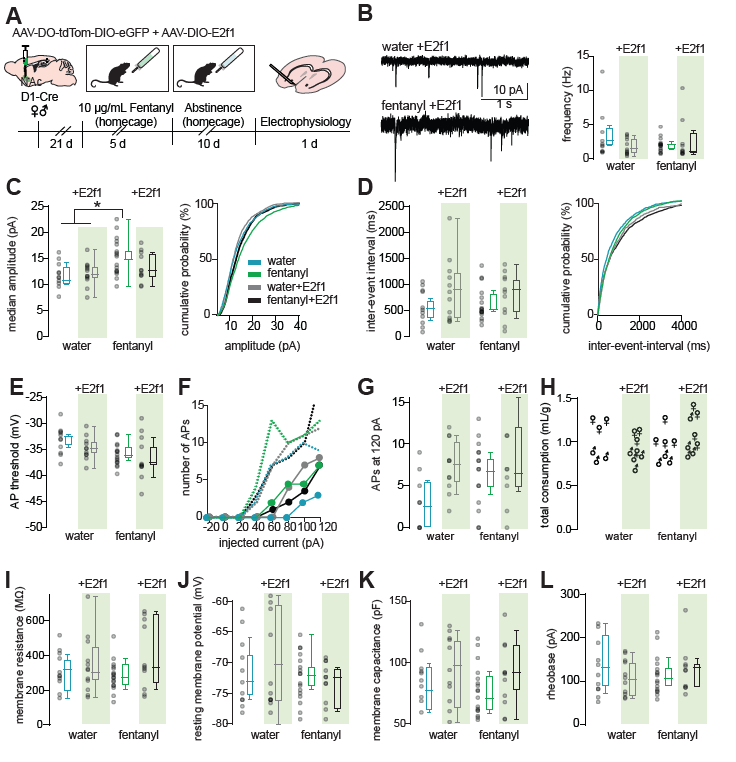
**Figure S5**

(**A**) Timeline for E2f1 electrophysiology experiments. Male and female D1-Cre mice received intra-NAc AAV-DO-TdTomato-DIO-eGFP to label MSN subtype and AAV-DIO-E2f1 for E2f1 overexpression in D1-MSNs. Following homecage water or fentanyl abstinence, slices containing the NAc were collected for patch-clamp recording. (**B**) Spontaneous excitatory postsynaptic currents (sEPSCs) were recorded while holding the membrane potential at -50 mV and analyzed by template-based event detection. Representative sEPSCs in water and fentanyl abstinent E2f1 expressing D1-Cre mice. Comparison of median sEPSC frequency between water and fentanyl abstinent mice expressing DO-tdTomato-DIO-eGFP with or without DIO-E2f1. (**C**) Comparison of median sEPSC amplitude (*, p’s<0.04 Fentanyl-GFP vs water-GFP, vs water-E2f1, Sidak’s after ANOVA), and cumulative probability in D1-MSNs from water and fentanyl abstinent mice that received DO-tdTomato-DIO-eGFP with or without DIO-E2f1(**D**) Comparison of median sEPSC inter-event-interval and cumulative probability in D1-MSNs from water and fentanyl abstinent mice that received DO-tdTomato-DIO-eGFP with or without DIO-E2f1. Individual data points represent individual neurons. Box and whiskers plots show the median and range for data nested by mouse. (**E**) Comparison of the membrane potential required to elicit an action potential (AP), and (**F**) number of APs elicited by injecting current in 20 pA intervals (median+range, circles and dashed lines, respectively), in D1-MSNs from water or fentanyl abstinent mice that received DO-tdTomato-DIO-eGFP with or without DIO-E2f1. (**G**) Comparison of the number of APs elicited by 120 pA injection in D1-MSNs from water and fentanyl abstinent mice that received DO-tdTomato-DIO-eGFP with or without DIO-E2f1. (**H**) Total liquid consumption across 5 days for male and female D1-Cre mice used in all electrophysiology experiments. Comparison of (**I**) membrane resistance, (**J**) resting membrane potential, (**K**) membrane capacitance, and (**L**) rheobase in D1-MSNs from water and fentanyl abstinent mice that received DO-tdTomato-DIO-eGFP with or without DIO-E2f1. Detailed Statistics in Supplemental File 1.

**Table S1**. **Related to Figure 3.**  WGCNA modules with significant effect of fentanyl abstinence. Module number and color, number of genes in the module, the top 10 hub genes, log fold change and P value for fentanyl abstinence in D1+D2, log fold change and P value in D2, log fold change and P value in D1

| **Module** | **Color** | **nGenes** | **top10 Hub Genes** | **logFC fentanyl** | **P.Value fentanyl** | **logFC fentanyl_in_d2** | **P.Value fentanyl_in_d2** | **logFC fentanyl_in_d1** | **P.Value fentanyl_in_d1** |
| --- | --- | --- | --- | --- | --- | --- | --- | --- | --- |
| D1.AE9 | blue | 206 | Dph1,Por,Rarb,Cpne2,Wdr18,Trap1,Mpp1,Ctnnbl1,Mgat2,Psen1 | 1.275507548 | 0.022152801 | 0.065283232 | 0.864442247 | 1.210224316 | 0.002687039 |
| D1.AE3 | gray | 417 | Arl6,Ptpra,Ndfip1,Vapa,Prps1,Psmc4,Serinc1,Nap1l2,Cops4,Ppp2ca | 1.058554651 | 0.011768987 | -0.024136538 | 0.932810796 | 1.082691189 | 0.000440705 |
| D1.AE11 | yellow | 104 | Prepl,Gm5499,Mtrex,Rars2,Ythdf2,Rnf180,Tm9sf2,Mmachc,Fbxl5,Fbxo3 | 0.765040912 | 0.028942308 | -0.179467198 | 0.457610027 | 0.944508109 | 0.000287229 |
| D2.AE14 | brown | 79 | Pdia6,Hspa5,Rexo4,Cys1,Dusp6,Egr4,Nr4a1,Atf3,Rnf32,Prps1l3 | 0.784637105 | 0.087862045 | -0.092543165 | 0.772253365 | 0.87718027 | 0.008416034 |
| D2.AE11 | purple | 155 | Bag5,Ptcd2,Fbxl5,Pitpnb,Timp2,Recql,Pcyox1,Rarb,Mgat2,Bhlhb9 | 0.604751646 | 0.185267545 | -0.152888763 | 0.632753305 | 0.757640409 | 0.021515141 |
| D2.AE8 | black | 194 | Tmem158,Hmbs,Asphd2,Angptl6,Xxylt1,Limd2,Rnf166,Ap1s1,Cnot8,Itpa | 0.754070936 | 0.044917604 | 0.42029745 | 0.110810545 | 0.333773485 | 0.202966648 |
| D2.AE2 | magenta | 361 | Tubb2b,Dlx1,Plpp3,Slc1a3,Arx,Fabp7,Pla2g7,Ndrg2,Mlc1,Cd9 | -0.979626559 | 0.001505807 | -0.817756422 | 0.00024248 | -0.161870138 | 0.432854847 |
| D2.AE15 | cyan | 72 | Ttc39b,Gria3,Kcna2,Csnk2a2,Bicd1,Ank2,Ppm1k,Cacna1a,Arap2,Map7d2 | -0.417589101 | 0.293221885 | 0.16685955 | 0.550809096 | -0.58444865 | 0.041023264 |
| D1.AE10 | red | 169 | Ctxn3,Trdn,Gcnt1,Nacc2,Gng4,Ntn4,Stard8,Kcnab3,Lgr6,Gprc5b | -1.385223155 | 0.006879069 | -0.498026518 | 0.156246406 | -0.887196637 | 0.013668682 |
| D1.AE4 | green | 388 | Cdk5r1,Syt7,Nlgn2,Bcl7a,Syngap1,Prkce,Nat8l,Rab35,Pura,Gramd1b | -0.971595699 | 0.026780641 | 0.023067331 | 0.939001119 | -0.99466303 | 0.001834251 |
| D1.AE18 | orange | 41 | Atf4,Ssh2,Ank2,Chmp4b,Dcp2,Lars2,Gm15564,Psmg3,Marf1,Pex11a | -1.267043609 | 0.010388206 | -0.027922498 | 0.933847195 | -1.239121111 | 0.000593379 |
