## Supplementary material for "Negative emotional behavior during fentanyl abstinence is mediated by adaptations in nucleus accumbens neuron subtypes": Key Resources Table

| REAGENT or RESOURCE | SOURCE | IDENTIFIER |
| --- | --- | --- |
| Antibodies | | |
| anti-E2f1 for western | Abcam, Waltham, MA, USA | Cat#ab137415 |
| anti-Prkce for western | Proteintech, Rosemont, IL, USA | Cat#20877-1-AP;RRID:AB_10697812 |
| anti-Gramd1b for western | Proteintech, Rosemont, IL, USA | Cat#24905-1-AP;RRID:AB_2879791 |
| anti-Nlgn2 for western | ProSci, Poway, CA, USA | Cat#7969 |
| anti-GAPDH for western | Cell Signalling, Danvers, MA, USA | Cat#2118;RRID:AB_561053 |
| Rabbit (DA1E) mAb IgG XP^®^ Isotype Control (CUT&RUN) | Cell Signalling, Danvers, MA, USA | Cat#66362 |
| Tri-Methyl-Histone H3 (Lys4) (C42D8) Rabbit mAb for Cut&Run | Cell Signalling, Danvers, MA, USA | Cat#9751;RRID:AB_2616028 |
| anti-E2f1 for Cut&Run | Invitrogen,Waltham, MA *USA* | Cat#32-1400;RRID:AB_2533065 |
| Bacterial and Virus Strains | | |
| AAV5-Ef1a-DIO-eYFP | UNC Vector Core, Chapel Hill, NC, USA | RRID:Addgene_27056 |
| AAV9-Ef1a-DO-TdTomato-DIO-EGFP | UMB Vector Core, Baltimore, MD, USA | RRID:Addgene_37120 |
| AAV9-Ef1a-DIO-E2f1-myc-Flag | This manuscript, UMB Vector Core, Baltimore, MD, USA | N/A |
| Chemicals, Peptides, and Recombinant Proteins | | |
| Fentanyl citrate | Cayman Chemical, Ann Arbor, MI, USA | Cat #22659 |
| Critical Commercial Assays | | |
| nCounter Master Kit--48 rxns NAA-AKIT-048 | Nanostring, Seattle, WA, USA | Cat#100054 |
| RNAscope® Probe - Mm-E2f1 | Advanced Cell Diagnostics,Newark, CA, USA | Cat#431971 |
| RNAscope® Probe - iCre-C2 | Advanced Cell Diagnostics,Newark, CA, USA | Cat#423321-C2 |
| RNAscope^®^ Fluorescent Multiplex Reagent Kit | Advanced Cell Diagnostics, Newark, CA, USA | Cat#320850 |
| Cut & Run kit | Cell Signalling, Danvers, MA, USA | Cat#86652 |
| nCounter GX CodeSet--48 rxns GXA-P1CS-048 | Nanostring, Seattle, WA, USA | Cat#100021 |
| Deposited Data | | |
| Differential expression and WGCNA | This paper | Mendeley Data, V1, doi: 10.17632/snpmrt8fj3.1 |
| RNA-seq data | This paper | GEO (TBD) |
| Experimental Models: Cell Lines | | |
| Neuro2A cells | Sigma | Cat#[89121404](https://www.sigmaaldrich.com/US/en/product/sigma/cb_89121404) |
| Experimental Models: Organisms/Strains | | |
| RiboTag mice;B6J.129(Cg)-*Rpl22^tm1.1Psam^*/SjJ | Bred at UMSOM | RRID:IMSR_JAX:029977 |
| A2A Cre mice;Tg(Adora2a-cre)KG139Gsat | Bred at UMSOM | Line KG139 |
| D1 Cre mice; Tg(Drd1-cre)FK150Gsat | Bred at UMSOM | Line FK150 |
| CD-1 mice, retired breeders | Charles River | RRID:IMSR_CRL:022 |
| C57Bl/6 mice | Bred at UMSOM | N/A |
| Oligonucleotides | | |
| Gramd1b Cut&Run primers | IDT DNA,  Redwood City, California, USA | Forward:TTAAGATGCGCCGGATGAAGA Reverse:GTCGCTGGAGGGCGTAATC |
| Nlgn2   Cut&Run primers | IDT DNA | Forward:AATCAGCATGTGGCTCCTGG Reverse:GATCTCGTTGTTGAGCTCGC |
| Prkce Cut&Run primers | IDT DNA | Forward:AGATCCGAGGAGCACAGACTC Reverse:ATAGAAAGTTTTGCCGGTTGGGG |
| E2F1 primers | IDT DNA | Forward:CTCGACTCCTCGCAGATCG Reverse:GATCCAGCCTCCGTTTCACC |
| Gramd1b  primers | IDT DNA | Forward:CTGCCGTCCATTGAGATTACG Reverse:TCAGGAACCTGCTCGAATCAT |
| Nlgn2 primers | IDT DNA | Forward:TGTCATGCTCAGCGCAGTAG Reverse:GGTTTCAAGCCTATGTGCAGAT |
| Prkce primers | IDT DNA | Forward:GGGGTGTCATAGGAAAACAGG Reverse:GACGCTGAACCGTTGGGAG |
| Recombinant DNA | | |
| Mouse E2f1-myc-Flag ORF clone | Origene | #MR206856 |
| Software and Algorithms | | |
| TopScan Lite | Cleversys, Reston, VA, USA | RRID:SCR_014494 |
| Imaris 8.3 | Bitplane, Oxford Intsruments | RRID:SCR_007370 |
| nSolver | Nanostring | RRID:SCR_003420 |
| pClamp | Axon Instruments | RRID:SCR_011323 |
| MiniAnalysis | Synaptosoft | RRID:SCR_002184 |
| TopHat |  | RRID:SCR_013035 |
| HTSeq |  | RRID:SCR_005514 |
| Weight Gene Co-Expression Network Analysis |  | RRID:SCR_003302 |
| BiNGO plug-in for Cytoscape |  | RRID:SCR_005736 |
